# Synthesis based on covalent capture and release (SCCR): a programmable strategy for automated preparation of protease-activatable molecules

**DOI:** 10.64898/2026.02.07.704608

**Authors:** Mayano Minoda, Tadahaya Mizuno, Takumi Iwasaka, Hiroyuki Kusuhara, Yu Kagami, Shingo Sakamoto, Norimichi Nagano, Chiaki Hori, Kazufumi Honda, Yasuteru Urano, Toru Komatsu

## Abstract

Enzyme-activatable chemical tools, including fluorogenic probes and prodrugs, are essential in chemical biology and targeted therapeutics but remain challenging to access in structurally diverse forms because their synthesis is often bespoke and difficult to standardize. Here, we introduce synthesis based on covalent capture and release (SCCR) as a programmable chemical strategy that enables the modular assembly of protease-activatable molecules through specifically designed protecting-group logic. The SCCR framework establishes a standardized capture–elongation– release workflow that decouples molecular diversification from individual synthetic optimization, thereby enabling automated preparation of complex libraries. Using this chemistry, we generated a diverse set of fluorogenic probes and applied them to single-molecule enzyme activity analyses to identify candidate activity-based biomarkers of liver diseases. The generality of the SCCR strategy was further demonstrated by extending the same chemical logic to the preparation of antibody–drug conjugate (ADC) linkers, allowing systematic evaluation of plasma stability and cytotoxic potential. By establishing a programmable capture–release chemistry for the synthesis of enzyme-activatable molecules, this work provides a generalizable chemical foundation for the scalable and automated construction of functional small-molecule tools across biological and translational research.

## Introduction

Enzyme-activatable chemical tools that exhibit specific functionalities upon targeted enzymatic reactions are powerful approaches for studying and controlling biological processes. To date, many chemical probes have been developed for use in biological, biochemical, and medicinal research and as therapeutic drugs. For example, fluorogenic probes are designed to fluoresce upon activation by the metabolism of a substrate, enabling studies of enzyme activity in complex biological systems^1,2^. Cancer-targeted prodrugs, such as antibody-drug conjugates (ADCs), exhibit cytotoxicity toward targeted tumor cells following activation by specific enzymes^3,4^.

Peptides and other modular molecular scaffolds represent a major platform for the design of chemical tools activatable by peptidases/proteases. As a variety of peptidases/proteases are present in living systems, both on-target and off-target effects must be considered to optimize probes to achieve desired functionality. An approach to develop such probes is to synthesize numerous candidates and screen them to identify those that exhibit the desired function in the biological systems of interest^5–7^. This requires the efficient synthesis of a diverse array of candidate probes in a high-throughput manner. However, conventional approaches for the synthesis of enzyme-activated chemical tools do not satisfy this demand.

The use of automated synthesis methods to generate candidate molecules for activity screening has increased in medicinal chemistry research^8,9^. However, these methods have not been fully exploited for the construction of enzyme-activatable chemical probe libraries. This is partially due to the difficulty of standardizing the purification steps of those chemical probes that have diverse molecular weights and charges^2^. In the present study, we report a programmable synthesis based on covalent capture and release (SCCR) strategy, in which a covalent capture handle is introduced into the protecting groups of the molecule and the desired product is purified by capture on a solid phase and release via deprotection. This approach can simplify and standardize the purification protocol, and therefore could lead to the fully automated development of enzyme-activatable chemical tools such as fluorogenic probes and prodrugs (**Figure 1a**).

**Figure 1.**
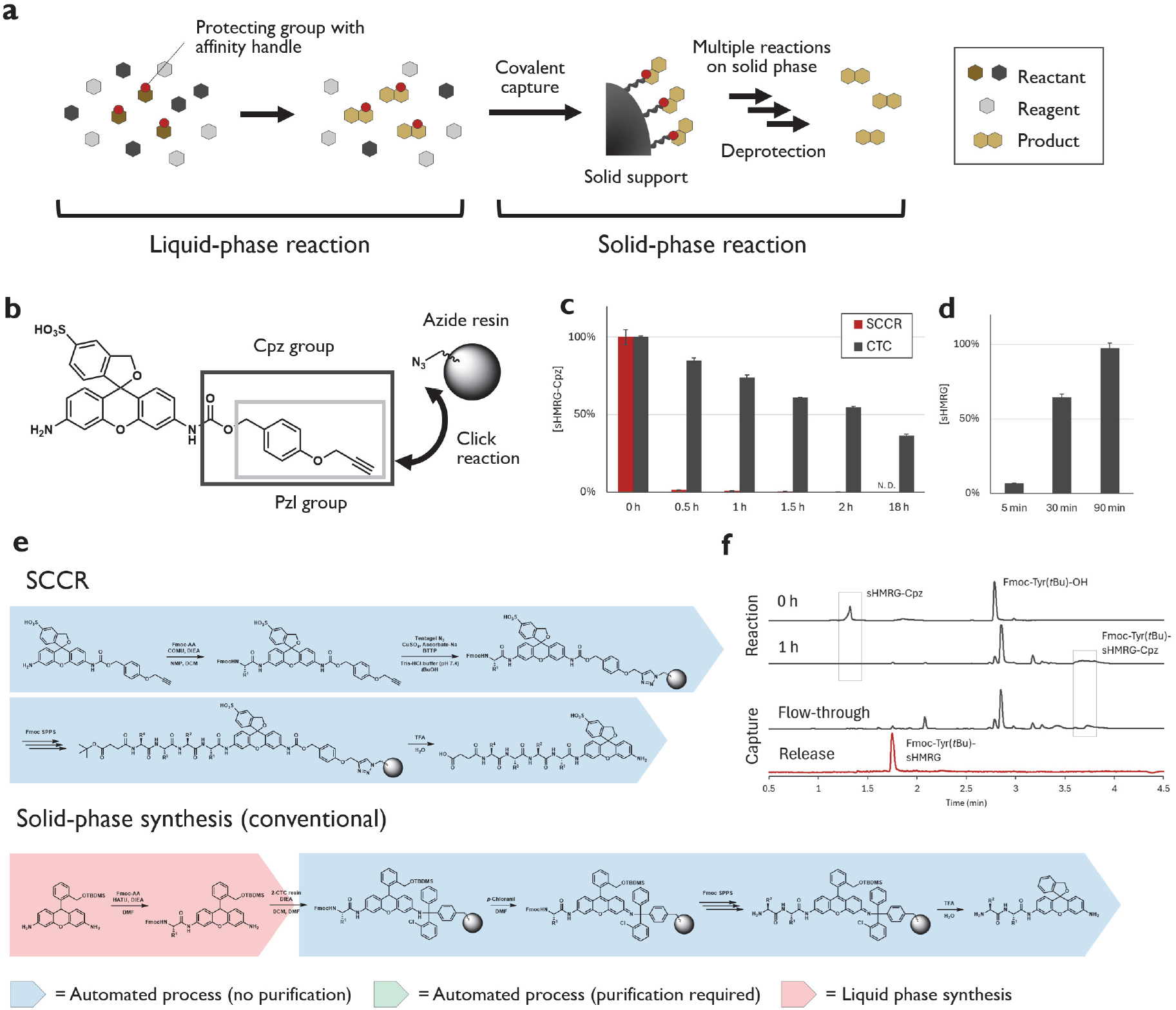
Concept and workflow of synthesis based on covalent capture and release (SCCR). (a) Conceptual workflow of SCCR by covalently capturable protecting groups. (b) Molecular structure of Cpz-modified sHMRG designed for SCCR using Tentagel-azide resin. (c) Capture of sHMRG-Cpz by azide resin (SCCR; red) and 2-chlorotrityl chloride resin (CTC; gray). For capture by azide resin, sHMRG-Cpz (0.5 μmol) was dissolved in 1 mL of Tris-HCl buffer (100 mM, pH 7.4) containing *t*-BuOH (40%), DMSO (10%), CuSO_4_ (1 mM), BTTP (3 mM), and sodium ascorbate (3 mM) and incubated with azide resin (0.26 mmol/g, 50 mg) at 25°C for the indicated times. For capture by CTC resin, sHMRG-Cpz (0.5 μmol) was dissolved in 400 μL of NMP and 100 μL of DIEA and incubated with CTC resin (1.55 mmol/g, 30 mg) at 25°C for the indicated times. The sHMRG-Cpz remaining in solution was quantified using LC-MS. Error bars represent standard deviation (S. D.; n = 3). N. D. = not detectable. (d) Release of azide resin–captured sHMRG-Cpz by acidic deprotection. For the beads captured under the conditions described in (c) for 2 h, beads were washed three times with DMF and DCM, and then 90% TFA-10% H_2_O (500 μL) was added, and the sample was shaken at 25°C for the indicated times. Released sHMRG was quantified using LC-MS. The recovery rate was calculated by normalization relative to sHMRG-Cpz (0.5 μmol) treated with 90% TFA-10% H_2_O (500 μL). Error bars represent S. D. (n = 3). (e) Scheme for preparation of sHMRG-based fluorogenic peptidase probes using SCCR. The representative conventional scheme for solid-phase synthesis-based fluorogenic probes^6^ is shown, and other schemes are summarized in **Figure S1.** (f) LC chromatograms (254 nm absorbance) monitoring the preparation of an sHMRG-based probe by attachment of Fmoc-Tyr(*t*Bu)-OH. The reaction was performed using a mixture of sHMRG-Cpz (1 μmol), DMT-MM (10 μmol), and Fmoc-Tyr(*t*Bu)-OH (10 μmol) in 20 μL of NMP, 10 μL of DIEA, and 500 μL of DCM with incubation at 55°C for 0 or 1 h (top chromatograms). The reaction product was dissolved in 400 μL of *t*-BuOH and added onto azide resin (0.26 mmol/g, 50 mg) dissolved in 300 μL of Tris-HCl buffer (100 mM, pH 7.4). Next, 100 μL of CuSO_4_ aq. (10 mM), 100 μL of BTTP solution in DMSO (30 mM), and 100 μL of sodium ascorbate aq. (30 mM) were added successively and incubated at 25°C for 2 h. The flow-through was then analyzed, and the beads were washed 6 times with 1.5 mL of DMF and 3 times with 1.5 mL of DCM. The beads were treated with 1 mL of 90% TFA-10% H_2_O for 10 min at 25°C, and the eluate was collected.

### Design of protecting groups for SCCR

Protease/peptidase activatable chemical probes have peptide backbones, and standard peptides can be prepared in an automated manner using Fmoc solid-phase peptide synthesis^10^. However, the process of attaching functional scaffolds, such as fluorophores or self-cleavable linkers, to generate activatable chemical tools is difficult to integrate into solid-phase synthesis schemes due to the limited scope of reaction conditions and low yields in solid-phase synthesis. For example, achieving a nearly 100% conversion rate in the functionalization of aniline to amide (anilide) using solid-phase synthesis is challenging, while these reactions are key steps in developing activatable chemical tools^5,6,11^. To overcome this limitation, we have proposed utilizing synthesis based on affinity separation (SAS)^5,12^, in which reactions are carried out in the liquid phase with reactants having an affinity handle, and products are purified using solid-phase affinity purification^5^. The system combines the advantages of liquid-phase reactions, which are suitable for a wide variety of reactions and provide good reaction yields, with those of solid-phase reactions, such as ease of purification (**Figure 1a, S1**). In the original SAS scheme, a handle for non-covalent affinity separation is attached to the probe scaffold^5^, but this imposes limitations on molecular design since the handle needs to be included in the final product. We considered that more flexible and versatile design becomes possible by SCCR through attaching the affinity handle to the protecting group and using covalent capture instead of non-covalent affinity purification. The design enables highly selective capture of the target molecule in a liquid-phase reaction mixture, enables additional reactions to be performed on a solid phase, and allows the release of the final product without the affinity handle upon deprotection of the protecting group.

### Automated synthesis of fluorogenic probes using SCCR

As the first application of this system, we established a method to generate a library of fluorogenic probes for reporting enzyme activity at the single-molecule level^2^. Sulfonated hydroxymethyl rhodamine green (sHMRG) is a fluorophore designed to specifically report the activity of peptidases/proteases in a microdevice^13,14^, and we attempted to make this fluorophore compatible with SCCR-based probe library preparation. We designed *p*-propargyloxybenzyl (Pzl) as an SCCR-compatible protecting group, based on the *p*-methoxybenzyl (PMB) group (**Figure 1b**). PMB is cleavable under acidic conditions such as trifluoroacetic acid (TFA) but stable under many other reaction conditions^15^. The handle, a propargyl group, can be captured using click chemistry with azide as a way to rapidly and selectively form a covalent bond. The Pzl group can be used to protect amines by forming a carbamate, and the carbamate-type protecting group was named p-propargyloxybenzyl carbamyl (Cpz), analogous to the Cbz group. As a capturing unit, PEG-lylated azide was modified on Tentagel-NH_2_ beads. In this synthetic scheme, a compound with a Pzl/Cpz group can be captured on a solid phase using click chemistry, and the final product without the protecting group can be selectively released from the beads using acidic deprotection.

Capture/release performance was evaluated using sHMRG-Cpz and compared with the performance of 2-chlorotrityl chloride (CTC) resin, which is commonly used in solid-phase synthesis. Capture using click chemistry was rapid, proceeding almost quantitatively within 15 min at a concentration of 0.1 M (**Figure 1c**). This was in sharp contrast with the modification of CTC resin, whose reaction with aniline was relatively slow and required an excess of reagents to fully load the resin with reactants. The captured Cpz group was stable, and cleavage and release were easily achieved using 90% TFA in H_2_O, a standard deprotection condition for acid-labile protecting groups. Almost quantitative recovery was achieved after 90 min (**Figure 1d**).

A key aspect of our scheme is the selectivity of the capture process, so the desired product in the liquid-phase reaction can be enriched on the solid phase for purification, successive reactions on the solid phase, and release by deprotection. Therefore, the inefficient anilide forming reaction with the first amino acid (P1) can be performed in the liquid phase^5^, and the subsequent purification process and peptide elongation steps can be performed on the solid phase (**Figure 1e, S1**). Almost quantitative conversion of aniline to anilide with an amino acid building block was realized by running the reaction (1) in the liquid phase, (2) with excess carboxylic acid and condensation reagents (5 eq.), (3) at high concentration (1 M), and (4) at high temperature (55-65°C)^5^ (**Figure 1f, top**). After the reaction, the product was captured using azide resin, the peptide was elongated on the solid phase, and the product was released from the resin. The product was of sufficient purity for a single-molecule enzyme activity assay (>90%, monitored at 500 nm) without requiring specific purification after the overall procedure (**Figure 1f**). Due to the simplicity of the scheme, we were able to automate the entire process using an automated peptide synthesizer. Compared with previous schemes employed to develop peptide-based fluorogenic probes^6^, the proposed scheme significantly reduced the number of processes requiring human manipulation (**Figure S1**). Using this scheme, we synthesized 103 probes that target various endo- and exo-peptidases (**Figure S2, S3, Table S1**). Peptidases were categorized as those targeting trypsin-like endopeptidases (P1 = Arg and Lys), chymotrypsin-like endopeptidases (P1 = Phe, Trp, and Tyr), caspase-like endopeptidases (P1 = Asp), elastase-like endopeptidases (P1 = Ala, Leu, and Val), prolyl endopeptidases (P1 = Pro), and SUMO-specific proteases (SENPs, P1 = Gly) (**Table S2**).

We next extended the generality of the assay by preparing probes for another fluorophore. Si-rhodamine is a fluorophore that emits fluorescence in the red region, making it useful for multi-colored, single-molecule enzyme activity profiling (SEAP, **Figure 2a**)^14,16^. We introduced two sulfonic acid moieties to generate disulfonated Si-rhodamine (dsSiR) and attempted to add an SCCR-compatible protecting group. The difference between dsSiR and sHMRG is that by protecting one of the rhodamine anilines using an amide/carbamate, sHMRG becomes a ring-closure (spirolactam) form, which renders the other aniline sufficiently reactive for amide formation, whereas dsSiR cannot become a spirolactam, leaving the other aniline unreactive (**Figure S4a**). The leuco-form of dsSiR was then prepared, and one aniline was protected using an alkyl-type SCCR-compatible protecting group (**Figure S4b**). We designed a trityl (Trt) group modified with an alkyne as the alkyl-type SCCR-compatible protecting group, which was designated ethynyl-Trt (Ert). The Ert group was attached to one of the amines of leuco-dsSiR to generate the building block for red-emitting probes (**Figure S1**). Probe synthesis was easily performed, just as in the case of sHMRG-Cpz, whereas one additional reaction, oxidation of the fluorophore after peptide elongation, could also be performed on the solid phase.

**Figure 2.**
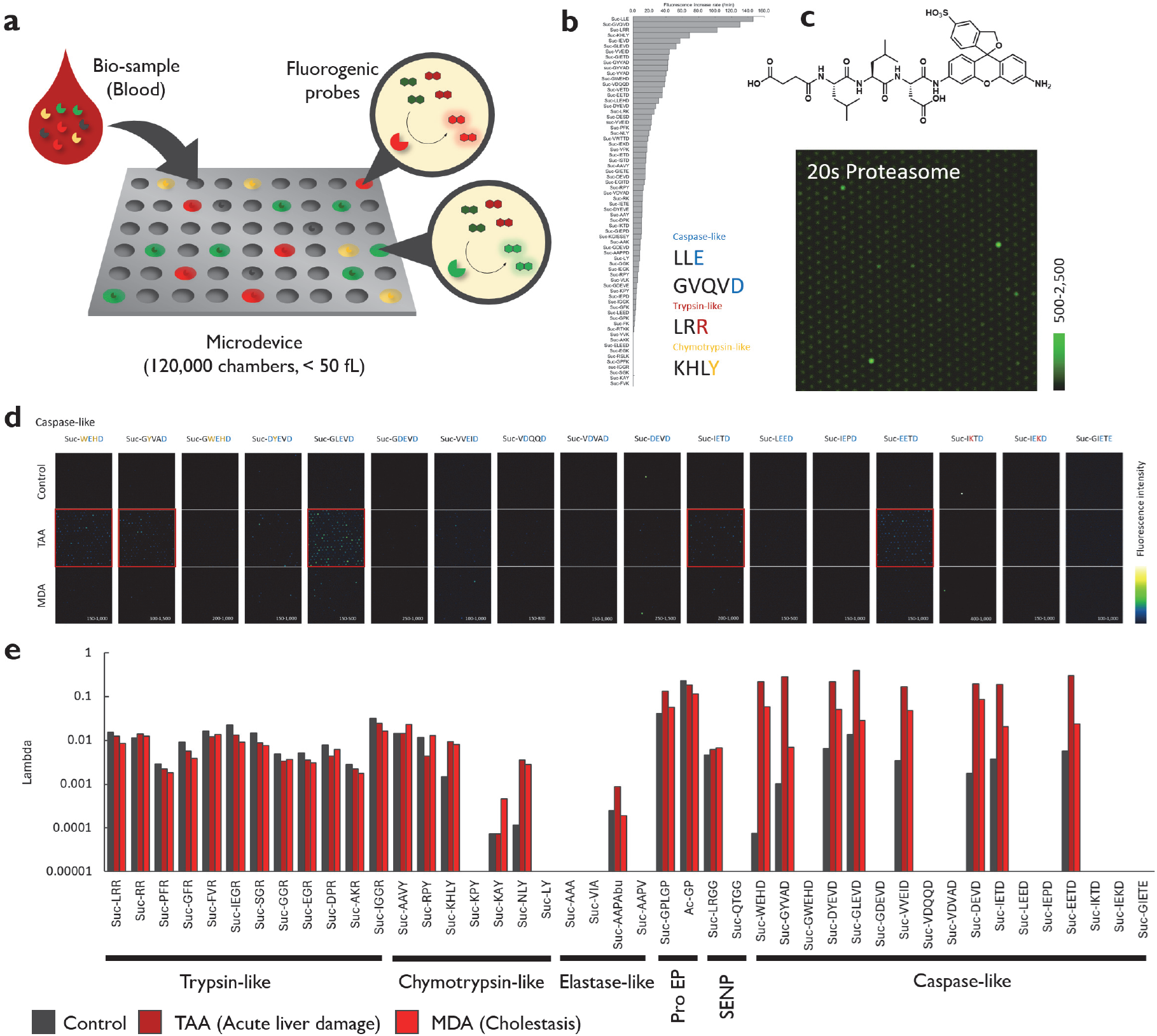
Establishment of activity-based screening to characterize single-molecule enzyme activities of recombinant enzymes and biomarker enzymes in blood samples (a) Concept of the single-molecule enzyme activity assay. (b) Substrate screening of recombinant proteasome 20S. Fluorogenic probes (10 μM) were mixed with proteasome 20S (activated by Triton X-100, 1 μg/mL) in HEPES-Na buffer (100 mM, pH 7.4) containing CHAPS (0.1%) and ATP (3 mM) and incubated for 60 min. The slope of the fluorescence increase was calculated. (c) Single-molecule enzyme activity assay of proteasome 20S. Suc-LLE-sHMRG (30 μM) was mixed with proteasome 20S (10 ng/mL) in HEPES-Na buffer (100 mM, pH 7.4) containing CaCl_2_ (1 mM), MgCl_2_ (1 mM), DTT (100 μM), and Triton X-100 (150 μM) and incubated at 25°C for 18 h. Fluorescence images were acquired using an epifluorescence microscope. (d) Result of activity-based screening of potential biomarkers of liver damage. The assay was performed by mixing probes (30 μM) with plasma samples (1/100, 1/1,000 for Pro-endopeptidases) in HEPES-Na buffer (100 mM, pH 7.4) containing CaCl_2_ (1 mM), MgCl_2_ (1 mM), DTT (1 mM), and Triton X-100 (250 μM) and incubating at 25°C for 18 h. Fluorescence images were acquired using probes for caspase-like activities. Imaging data for all probes are shown in **Figure S5.** (e) The number of positive spots observed in the experiments described in (d).

### Substrate profiling of recombinant proteasome 20S using single-molecule enzyme activity assays

With a library of 103 fluorogenic probes in hand, we first sought to identify substrates capable of reporting proteasome activity at the single-molecule level. Single-molecule enzyme activity assays were performed by introducing enzyme solutions into arrays of confined microreactors, where enzymatic turnover was detected by the accumulation of fluorescent reaction products (**Figure 2a**)^5,13,14,17–19^. While many single-molecule enzyme activities are becoming amenable to monitoring using this platform^2^, single-molecule assays for reporting proteasome activity have not been reported. The proteasome is a large multi-subunit protease complex responsible for regulated protein degradation and plays essential roles in diverse cellular processes, including protein quality control and antigen presentation^20^. The 20S core particle possesses multiple catalytic activities, typically classified as caspase-like, trypsin-like, and chymotrypsin-like activities, and their activities are subject to various posttranslational modifications^20–22^. To evaluate whether the SCCR-enabled probe library can identify suitable fluorogenic probes to study proteasome activity at the single-molecule level, we performed substrate profiling using recombinant human proteasome 20S. We identified multiple substrates that were metabolized by the proteasome (**Figure 2b**). The hit probes included known proteasome substrate sequences, such as Leu–Leu–Glu (LLE) and Leu–Arg–Arg (LRR), which preferentially report caspase-like and trypsin-like activities, respectively^21^. In addition, the profiling revealed previously unrecognized sequences, including Lys–His–Leu–Tyr (KHLY), which exhibited higher chymotrypsin-like activity than the commonly used Ala–Ala–Val–Tyr (AAVY) motif, and Gly–Val–Gln–Val–Asp (GVQVD), a sequence originally developed for caspase-3 and -6, that also reported caspase-like proteasome activity. The best substrate among them was LLE, which successfully reported proteasome activity on the single-molecule enzyme activity assay platform (**Figure 2c**). These results establish the feasibility of combining SCCR-based probe synthesis with single-molecule assays for profiling complex protease activities. This validation in a defined recombinant system motivated us to apply the same probe library to complex biological samples, where enzyme activities cannot be attributed to a single enzyme species but instead emerge as composite activity patterns.

### Activity-based screening to identify specific biomarker activities of liver damage

Having established that the SCCR-enabled probe library can identify suitable substrates and enable single-molecule measurements in a defined recombinant system, we next asked whether the same probes could extract diagnostically meaningful enzyme activity patterns from complex biological samples. In contrast to the recombinant protein system, in complex biological systems, a single fluorogenic probe can act on multiple enzymes and proteoforms with distinct activities. Therefore, candidate biomarkers are identified by analyzing the activity profiles obtained from each probe and searching for unique alterations in the patterns of single-molecule enzyme activities^2^. As a proof-of-concept, we searched for the blood-based biomarkers of liver damage to identify candidate biomarkers of liver disease. Two mouse models of liver damage with distinct molecular mechanisms were examined. The thioacetamide (TAA) model reflects hepatic cell damage^18^, whereas the 4,4’-methylene dianiline (MDA) model reflects cholestasis^23^. The current biomarker, aspartate/alanine aminotransferase (AST/ALT), cannot discriminate between these two liver diseases^24^, but we hypothesized that monitoring unique changes in enzyme activity patterns could distinguish the different pathologies.

Screening was performed using 44 representative probes selected from a library of 103 probes. Some of the probes showed notable differences in enzyme activities between control (vehicle-treated), TAA-treated, and MDA-treated mice (**Figure 2d, 2e, S5**). In TAA-treated mice, enzymes responding to Glu-Leu-Glu-Val-Asp (GLEVD)-sHMRG were upregulated (**Figure 3a, 3b**). The sequence is a substrate of caspases, and other probes targeting caspases also showed increased spot numbers in TAA-treated mice (**Figure 2d**). TAA damages hepatic cells^24^, and these results thus indicated the occurrence of apoptosis in liver tissues, resulting in leakage of caspases into the circulation. In MDA-treated mice, enzymes responding to Leu-His-Leu-Tyr (LHLY)-sHMRG (targeting kallikreins) were upregulated (**Figures 2e, S5**). The sequence targets multiple subtypes of kallikreins and other chymotrypsin-like proteases, and better dissection of the activity species was desired to characterize the molecular species contributing to the increase. To address this limitation, we introduced a second fluorogenic probe with a distinct color to enable multi-colored SEAP assays (**Figure 2a**). In the MDA model, the probe pair consisting of Suc-LHLY-sHMRG (green) and Suc-AAVY-dsSiR (red) revealed significant increases of enzyme spots exhibiting high activity toward both probes (**Figure 3c, 3d, S6**). Treatment with MDA triggers the activation of various immune cells accompanying cholestasis^24^; thus, we considered that specific kallikrein species are upregulated in response to the activation of these cells. Although a complete understanding of the molecular background will require further investigations, these results demonstrate the power of the proposed screening strategy using a panel of single-molecule enzyme assays to identify biomarker enzymes that reflect the pathological states of cells and tissues.

**Figure 3.**
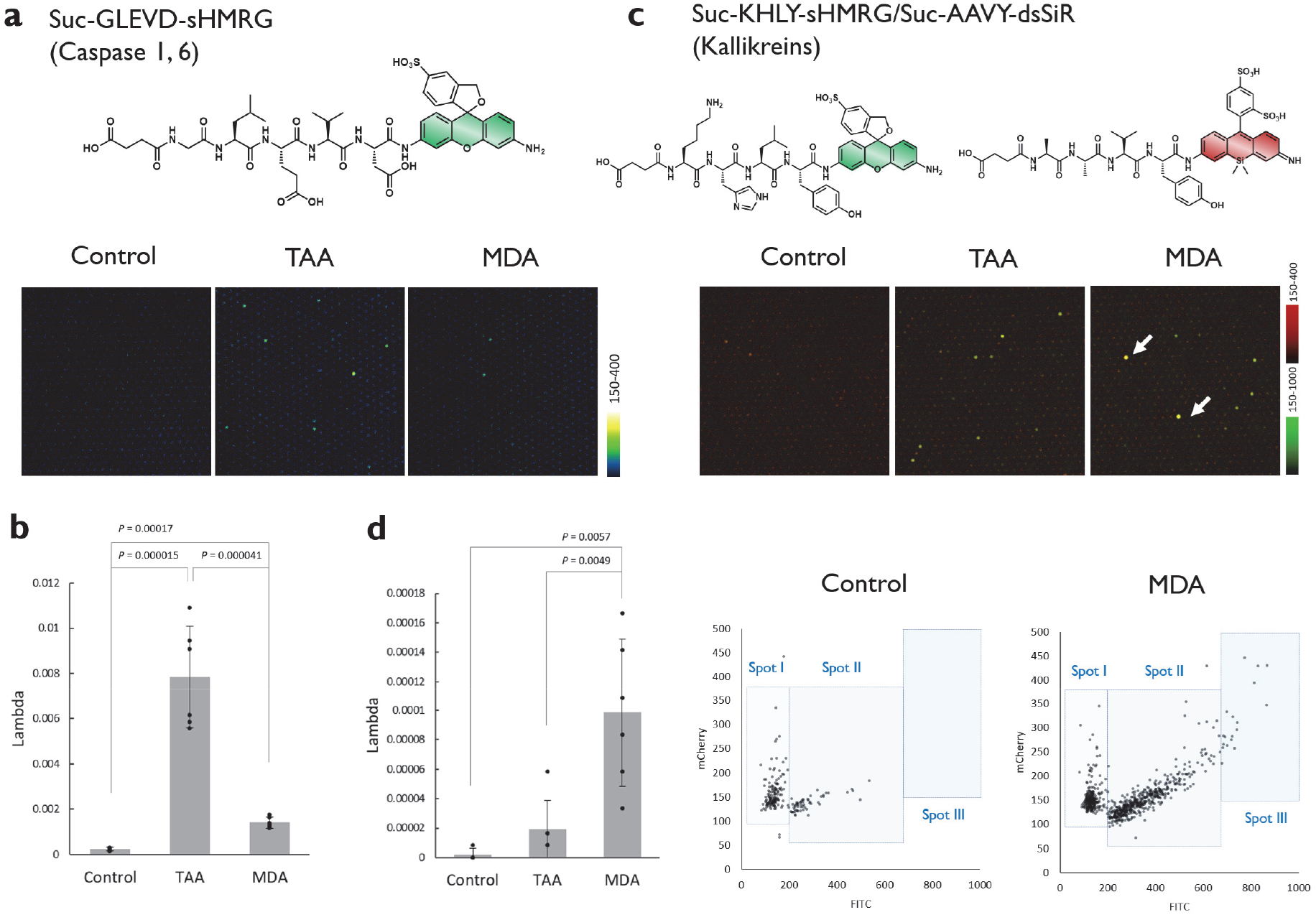
Identification of single-molecule enzyme activity-based biomarkers of liver damage with distinct mechanisms. (a) Fluorescence images of the microdevice after loading the probe (Suc-GLEVD-sHMRG) and blood samples. The assay was performed by mixing the probe (30 μM) with plasma samples (1/3,000) in HEPES-Na buffer (100 mM, pH 7.4) containing CaCl_2_ (1 mM), MgCl_2_ (1 mM), DTT (1 mM), and Triton X-100 (250 μM) and incubating for 18 h at 25°C. (b) Quantification of bright spots in (a); n = 4 for control and n = 6 for TAA- or MDA-treated mice. Error bars represent S. D. (c) Fluorescence images of the microdevice after loading the probe (Suc-GLEVD-sHMRG) and blood samples. The assay was performed by mixing the probe (30 μM) with plasma samples (1/100) in HEPES-Na buffer (100 mM, pH 7.4) containing CaCl_2_ (1 mM), MgCl_2_ (1 mM), DTT (1 mM), and Triton X-100 (250 μM) and incubating for 18 h at 25°C. (d) Scatter plots of activity spots shown in (c) with green fluorescence (sHMRG; horizontal axis) and red fluorescence (dsSiR; vertical axis). Quantification of spots observed in the area of spot III was compared using dot plots; n = 4 for control and n = 6 for TAA- or MDA-treated mice. Error bars represent S. D.

### Extension of the SCCR scheme to automated synthesis of prodrugs activated by specific proteases

We extended the concept of SCCR to synthesize a panel of ADC linkers using automated synthesis. ADCs have emerged as a promising therapeutic modality in cancer treatment^3,4^. The key to developing good therapeutics is to ensure the selection of a proper combination of antibody and ADC linkers with the payload; suitable ADC linkers also need to be stable in the blood in order to prevent non-specific toxicity^25^. However, the need for multiple synthetic steps in the synthesis of ADC linkers hampers diversity-oriented synthesis of candidate compounds (**Figure 4a, 4b, S7**).

**Figure 4.**
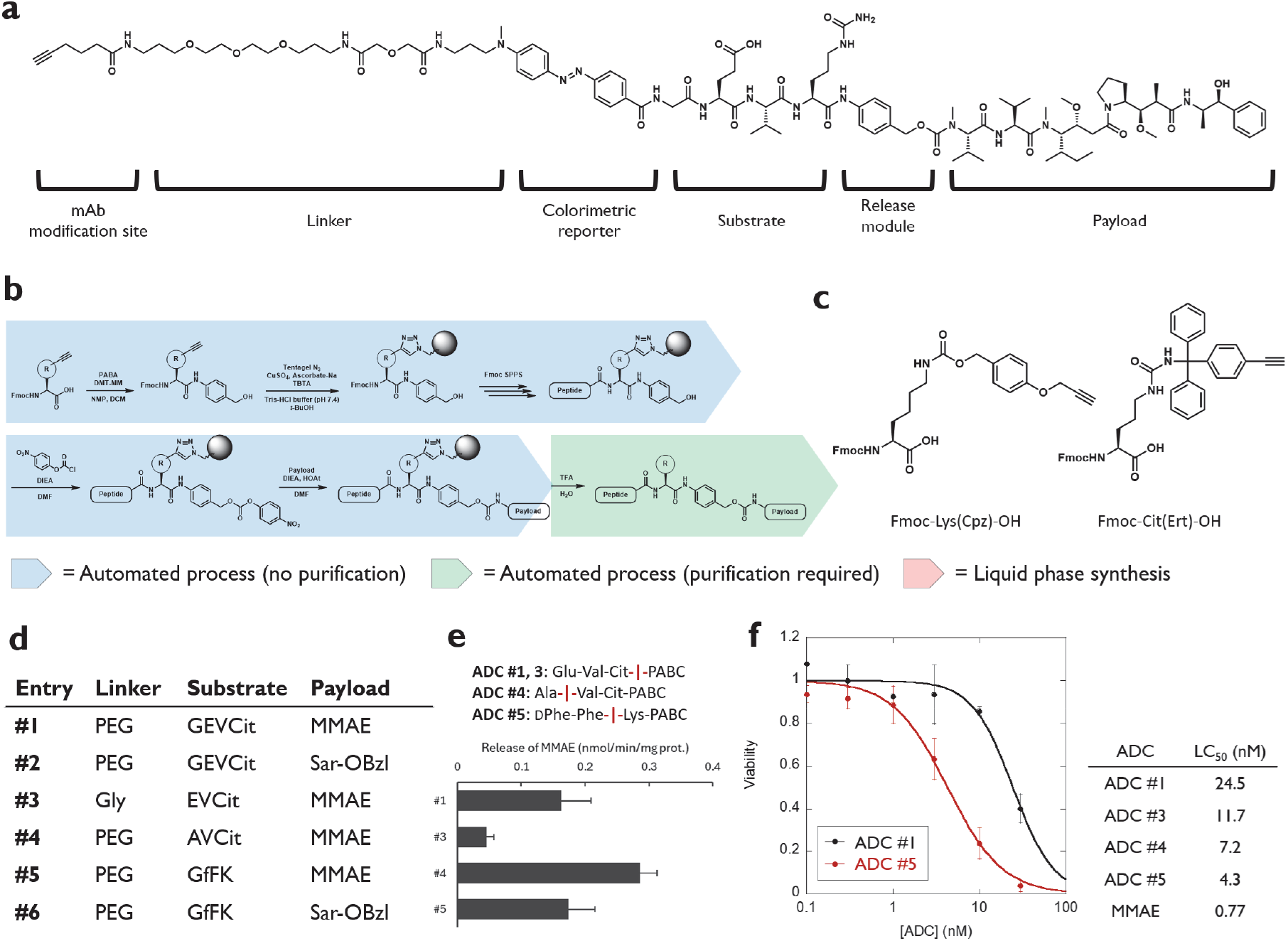
Automated synthesis of ADC linkers. (a) Structure of an ADC linker prepared in this study. (b) SCCR scheme for the preparation of ADC linkers. (c) Structures of amino acid derivatives for SCCR. (d) Panel of ADC linkers prepared in this study. The linker, substrate, and payload correspond to the structure shown in (a). PEG corresponds to NH_2_-PEG_3_-DGA-OH, and Sar-OBzl corresponds to *N*-methylglycine (sarcosine) benzyl ester. (e) Formation of MMAE after incubating ADC linkers (#1, #3, #4, and #5, 10 μM) with plasma samples obtained from healthy human subjects (1/4 dilution) in phosphate buffer (100 mM, pH 7.4) containing CHAPS (0.1%) for 3 h. Error bars represent S. D. (n = 3). The major cleavage site determined in mouse plasma samples is shown. Data for identification of the cleavage sites in mouse plasma are shown in **Figure S10.** (f) LC_50_ values against SKBR3 cells of trastuzumab modified with various ADC linkers. Detailed results are shown in the Supplementary Information. Error bars represent S. D. (n = 3).

We extended the concept of protecting group–based SCCR by attaching the protecting group to the amino acid building blocks introduced in the initial step of synthesis (**Figure 4b**). The initial step of the conventional ADC linker synthesis scheme involves condensation of *p*-aminobenzyl alcohol (PABA) with an amino acid building block (**Figure S7**). During synthesis, both functional groups of PABA, the aniline and the alcohol, are functionalized, so they cannot accept SCCR-protecting groups. However, we realized SCCR by attaching the SCCR-compatible protecting group to the side chains of the amino acids (**Figure 4b, 4c**). Twelve of the 20 standard amino acids used as building blocks have acid-labile protecting groups on their side chains, making them suitable for accepting SCCR-compatible protecting groups. As a representative example, we prepared a building block consisting of Lys and citrulline (Cit). With regard to Lys, the side chain amine was protected by a Cpz group. *N*-Cpz-protected Lys was stable under peptide elongation conditions (e.g., 40% *N*,*N*-diisopropyl-*N*-ethylamine (DIEA) in *N*,*N*-dimethylformamide (DMF) or 40% piperidine in *N*-methylpyrrolidone (NMP)), but deprotection was easily achieved under acidic conditions (e.g., 90% TFA-10% H_2_O), as for the Boc group (**Figure S8**). Similarly, the urea of Cit was successfully protected using an Ert group (**Figure S9**). Using these SCCR-compatible building blocks, we prepared ADC linkers with the sequence Xaa-Val-Cit and D-Phe-Phe-Lys. The former is a well-established sequence of ADC linkers that target various cathepsins^25^, and the latter is derived from a prodrug reportedly selective for cathepsin B^26^. The molecules included a chromophore-containing amino acid (3-aminopropyl-dabcyl; Apd)^17^ for ease of detection and quantification. The antibody attachment site was an alkyne moiety, and the payload was MMAE (monomethyl auristatin E) or *N*-methylglycine (sarcosine) benzyl ester (Sar-OBzl, control drug); linkers of differing lengths were included using polyethylene glycol (PEG) or glycine. We were able to prepare a panel of ADC linkers via automated synthesis, with purification using liquid chromatography required only at the final step (**Figure 4b**).

We generated six derivatives with varied structural features. First, we studied the stability of the derivatives in plasma obtained from healthy human subjects (**Figure 4d, 4e**). Stability in blood varied between the derivatives, with differences observed even between molecules bearing the same recognition sequences but different linker structures (Glu-Val-Cit sequence with a PEG linker [#1] and a glycine linker [#3]). With regard to differences in stability with different recognition sequences, cleavage at amino acids surrounding the desired cleavage site appeared to be problematic. We therefore characterized the cleavage sites of the ADC linkers. Since mouse plasma showed higher activity compared with human plasma, mouse plasma was therefore used for product analysis (**Figure S10**). LC-MS analysis indicated that ADC linkers with Glu-Val-Cit (#1-#3) were primarily cleaved after Cit, whereas the Ala-Val-Cit recognition sequence (#4) was cleaved at Ala-|-Val, and this undesired cleavage likely contributed to the instability of this linker in blood^25^. D-Phe-Phe-Lys (#5) was also cleaved at an undesired site, Phe-|-Lys, likely by proteases with chymotrypsin-like activity, but the activity was not high compared with that of Ala-Val-Cit (**Figure 4e**). These results highlight the importance of considering the activity of off-target enzymes in designing the recognition sequences of selective ADC linkers. The synthesized linkers were subsequently conjugated to the anti-HER2 antibody trastuzumab, and the resulting ADCs were evaluated for their cytotoxic activity against HER2-overexpressing SKBR3 breast cancer cells. All MMAE-bearing ADCs (#1, #3, #4, and #5) exhibited dose-dependent cytotoxicity, whereas the control conjugates lacking the payload (#2 and #6) showed no discernible effect (**Figures 4f, S11a**). The ADC incorporating the D-Phe-Phe-Lys recognition motif (#5) displayed distinct and notable activity in this cellular assay. Although the Ala-Val-Cit-based conjugate (#4) also exhibited high cytotoxicity, its activity was not suppressed by competition with excess trastuzumab, suggesting that its effect likely stemmed from non-specific linker cleavage within the serum-supplemented culture medium rather than target-specific internalization (**Figure S11a**). These observations underscore that the biological efficacy of an ADC is governed by a complex interplay of parameters, making it challenging to predict cellular performance solely based on i*n vitro* linker reactivity. Indeed, while the MMAE-releasing activity in SKBR3 cell lysates was comparable across conjugates #1, #3, #4, and #5 (**Figure S11b**), their actual cellular impacts varied significantly. Although further optimization is necessary to validate these leads in *in vivo* settings, these results highlight the utility of the SCCR platform in enabling the rapid preparation of structurally diverse candidate libraries, thereby facilitating empirical *in cellulo* screening required to identify functionally optimized linkers with the requisite balance of stability and reactivity.

## Discussion

In this study, we propose the concept of a programmable synthesis based on covalent capture and release (SCCR), in which a protecting group is repurposed as a covalent handle for purification, and demonstrate the utility of this approach for the automated synthesis of fluorogenic probes and ADC linkers. In the original synthesis based on affinity separation (SAS) strategy, the affinity handle was attached directly to the probe scaffold^5^. This design limited structural versatility because the handle had to remain in the final product, and the non-covalent nature of the interaction restricted purification to relatively mild conditions (**Figure S1**). We overcame these limitations by introducing the affinity handle into the protecting group itself. SCCR-compatible protecting groups behave like conventional protecting groups during liquid-phase synthesis, while enabling selective covalent capture of the desired intermediates on a solid phase. Subsequent reactions that tolerate excess reagents and proceed with near-quantitative yields can then be performed on the solid phase, followed by release of the final product through standard deprotection. The ease of standardization and the flexibility in designing synthetic routes render the SCCR scheme particularly well suited for fully automated chemical synthesis.

Screening enzymatic activities using an enzymomics-based approach has proven useful for discovering activity-based biomarkers^6,7,28^, in which libraries of enzyme substrates are prepared and screened against biological samples to identify activities of interest^6,7,29–32^. In this work, we developed a platform for screening single-molecule enzyme activities in biological systems and identified activity-based biomarkers capable of discriminating between distinct liver disease models. In parallel, the generality of the SCCR scheme was extended to the preparation of ADC linkers with diverse structures. Although purification was still required at the final step, primarily due to the unoptimized carbamate-forming reaction used to attach the payload (MMAE) and the acid lability of carbamates, the overall purification burden was substantially reduced compared with conventional ADC linker synthesis schemes^25^. Besides increasing synthetic throughput, the ability to perform many steps without direct human intervention offers an important practical advantage by reducing exposure to highly toxic payloads such as MMAE. Using this platform, we prepared a panel of ADC linkers and evaluated their stability in blood and cytotoxicity toward tumor cells. These analyses revealed that relatively subtle changes in linker length or amino acid sequence can markedly affect both stability and biological activity, underscoring the importance of programmable and scalable synthetic platforms.

In the present study, peptide sequences of chemical tools were designed based on existing knowledge of preferred substrate motifs for target enzymes (**Table S1**). The impact of automated synthesis platforms such as SCCR could be further amplified by integration with artificial intelligence (AI)–driven de novo molecular design, in which candidate compounds are generated based on bioactivity data from training sets^33,34^. By enabling rapid and systematic generation of diverse chemical libraries, the platform described here could serve as a powerful source of high-quality data for advancing data-driven chemical probe and drug design.

## Conclusion

We establish SCCR as a programmable synthetic strategy, in which a click reaction–based handle is incorporated into a protecting group rather than the core molecular scaffold. This fundamental shift in design enables substantial flexibility in both final product architecture and synthetic route selection. Ideally suited for fully automated synthesis, this approach facilitates the rapid preparation of diverse panels of chemical tools, including fluorogenic probes for single-molecule enzymomics and ADC linkers for high-throughput screening in biological systems. Using this platform, we identified candidate blood biomarkers associated with liver damage and discovered functionally optimized ADC linker sequences with tailored stability and potency for targeting breast tumor cells.

By streamlining the construction of sophisticated molecules through a standardized and reproducible workflow, SCCR significantly lowers the technical barriers traditionally associated with complex chemical synthesis. Ultimately, this platform empowers a broader range of researchers across the life sciences to independently develop custom chemical tools, fostering a more direct and efficient integration of synthetic chemistry into the heart of biomedical discovery.

## ASSOCIATED CONTENT

### Supporting Information

Methods and supporting tables/figures.

## AUTHOR INFORMATION

### Competing Financial Interests

T. M., K. H., and T. K. are advisors and shareholders of Cosomil, Inc. M. M., Y. U., and T. K. are inventors on patents covering the synthetic schemes and compounds reported in the article.

## Acknowledgements

This work was financially supported by MEXT (20H04694, 21A303, 22H02217, 23K23484, 25K01911, and 25K22520 to T. K.; 21K06663 to T. M.; 25H00241 and 25K23766 to N. N.), JST (PRESTO [13414915], PRESTO Network [17949814], CREST [19204926], and FOREST [24012649] to T. K.; START [20353017] to T. M., K. H., and T. K.), and AMED (FORCE [22581634] to T. M., K. H., and T. K.; P-PROMOTE [25131640] to T. M., K. H., and T. K.; Research on Development of New Drugs [23809006] to T. M. and T. K.; P-PROMOTE [18cm0106403h0003] to K. H.; P-CREATE [25ama221431h0002] to K. H.). T. K. received support from the Naito Foundation, The Mochida Memorial Foundation for Medical and Pharmaceutical Research, the Chugai Foundation for Innovative Drug Discovery Science, the MSD Life Science Foundation, the Hoansha Foundation, and the University of Tokyo Gap Fund Program.

