## Supplementary_Information for "Synthesis based on covalent capture and release (SCCR): a programmable strategy for automated preparation of protease-activatable molecules"

**Contents**

**Methods**

**Supplementary table S1-S2**

**Supplementary figure S1-S11**

**Supplementary methods for synthesis and characterization of compounds**

**Supplementary references**

### Methods

#### Materials

Reagents and solvents were of the best grade available, and were supplied by Tokyo Chemical Industries, Wako Pure Chemical, Sigma-Aldrich, Dojindo, Kanto Chemical Co., and Watanabe Chemical Industries, and were used without further purification. Enzymes were purchased from Sigma-Aldrich and were used without further purification.

#### Instruments

NMR spectra were recorded on a JEOL JNM-LA400 instrument at 400 MHz for $^{1}$H NMR and at 100 MHz for $^{13}$C NMR. Mass spectra (MS) were measured with a JEOL JMS-T100LC AccuToF (ESI). LC-MS analyses were performed on a Waters Acquity UPLC (H Class)/QDa quadrupole MS analyzer or Acquity UPLC (H Class)/Xevo TQD quadrupole MS/MS analyzer equipped with an Acquity UPLC BEH C18 column (Waters). Column chromatography using silica gel was performed on an MPLC system (Yamazen Smart Flash EPCLC AI-5805 (Tokyo, Japan)). Reversed-phase MPLC purification was performed on an Isolera One (Biotage) equipped with a SNAP Ultra C18 30 g (Biotage).

#### Activation of proteasome 20S

Recombinant proteasome 20S (R&D Biosystems, E-360, Lot #DBGH0724071) was activated by incubating the protein (10 μg/mL) in HEPES-Na buffer (100 mM, pH 7.4) containing Triton X-100 (0.03%) at 25°C for 30 min. The enzyme was diluted to the indicated concentrations for assays.

#### Digital enzyme assay in microdevice

Digital enzyme assays were performed using commercially available microdevices[1] (Simoa disk; Quanterix). A 40 μL of mixture of enzyme and reagents in buffer was loaded into the microdevice by manual pipetting. Then, 80 µL of FC-70 (Sigma-Aldrich) was introduced into the device to flush out excess reaction mixture. The enzymatic activity in the chambers was measured using an epifluorescence microscope (Ti2, Nikon) equipped with a 20× dry objective lens (Plan Apo 20×), an sCMOS camera (ORCA-Fusion C14440, Hamamatsu Photonics), a white LED illumination unit (X-Cite Xylis, Opto Science), and a motorized stage. The sHMRG-based assay was performed using a solution containing dsSiR (10 µM) as the internal standard, and the focus was adjusted using its fluorescence. Images were acquired in tile scan mode with perfect focus. The excitation and emission filters used were FITC (mirror = 510 nm, Ex. = 460-500 nm, Em. = 510-560 nm) and mCherry (mirror = 600 nm, Ex. = 550-590 nm, Em. = 608-683 nm), respectively.

**Image processing**

Images were processed using the GA3 module of NIS Elements software (Nikon). First, all fluorescence images were background-corrected using a rolling ball correction (3 µm). Then, ROIs were chosen by bright spot detection using mCherry filter settings (diameter = 3 µm, contrast = 500), and irregular fluorescent spots derived from fluorescent debris or air bubbles were omitted by dilating the ROI and removing the overlapping ROIs. The fluorescence signals were calculated as the mean of the signals from the center of each ROI. Data were processed using Excel, or Kaleidagraph software to construct histograms and scatter plots.

**Plasma samples from healthy human subject**

Plasma samples were collected from Kagoshima Prefectural Comprehensive Health Center with the Program for Promotion of Fundamental Studies in Health Sciences conducted by the National Institute of Biomedical Innovation of Japan, Health and Labour Sciences Research Grants from the Ministry of Health, Labour and Welfare of Japan, and P-CREATE of the Japan Agency for Medical Research and Development (AMED)36–39. Ethical approval for this study was obtained from the central ethics committee of Nippon Medical School (M-2021-002) and the ethical committee of Nippon Medical School (A-2020-032 and A-2020-044).

**Plasma samples from mice**

Ethical approval for the study using animals was obtained from the Animal Care and Use Committee of The University of Tokyo (P4-21, P31-9). Six-week-old male C57BL/6JJcl mice were purchased from CLEA Japan (Tokyo, Japan) and acclimatized for five days. The mice were exposed to thioacetamide (TAA, T0817, Tokyo Chemical Industry Co., Japan, 350 mg/L) or 4,4'-methylene dianiline (MDA, M0220, Tokyo Chemical Industry Co., Japan, 750 mg/L) dissolved in drinking water to induce liver damage, whereas control mice received tap water. After four days of treatment, mice were euthanized and blood was collected from the inferior vena cava into 1.5 mL tubes containing 1.5 µL heparin (Yoshindo Inc, Japan). The collected blood sample was centrifuged (1,700 g, 4°C for 15 min) for plasma separation. Characterization of the blood samples and the detailed disease states of mice are shown in the cited reference[2,3].

**Cell culture**

SKBR3 cells were cultured in 10 cm culture dishes. The culture medium was RPMI1640 (RPMI1640, Gibco), supplemented with 10% (v/v) fetal bovine serum and 1% (v/v) penicillin-streptomycin. Cells were cultured in a humidified incubator at 37°C under 5% $CO_2$ in 95% air.

**CCK-8 assay**

Cells were plated onto sterile clear 96-well plates (Nunc 267334) at 1,000 cells/well. Varied concentrations of trastuzumab modified with ADC linkers were added to wells and cells were cultured for 12 h. For competition conditions, ADCs were co-incubated with trastuzumab (100 nM). Media were removed and 200 µL resh media was added, and cells were cultured for 48 h. Media was removed and 100 µL media containing 10 µL CCK-8 solution (Dojindo CK04) was added. After incubating for 2 h, absorbance at 450 nm was measured using EnVision. For calculating 100% viability, wells without treatment were used, and for 0% viability, wells with cells treated with 70% EtOH were used.

**Preparation of cell lysate**

SKBR3 cells were cultured in 10 cm culture dishes to 50-60% confluency. The cells were washed twice with PBS, and 500 µL CelLytic M was added. After gently shaking at 25°C for 10 min, the lysate was collected and centrifuged (14,000 rpm × 10 min, 4°C). The supernatant was collected, and aliquoted for single use. Protein concentration was determined by Bradford assay using bovine serum albumin (BSA) as a standard.

**Metabolism of ADC linkers in biological samples**

For analysis of MMAE formation from ADC linkers in plasma samples, ADC linkers (10 µM) were mixed with plasma samples from healthy human subjects (1/4 dilution) in phosphate buffer (100 mM, pH 7.4) containing CHAPS (0.1%) and incubated at 37°C for 3 h. For analysis of MMAE formation from ADC linkers in cell lysate, ADC linkers (30 µM) were mixed with lysate of SKBR3 cells (0.15 mg/mL) in phosphate buffer (100 mM, pH 5.5) containing DTT (1 mM) and CHAPS (0.1%) and incubated at 37°C for 20 h. The reaction was mixed with an equal volume of AcCN-10% formic acid. After centrifugation (14,000 rpm × 10 min, 4°C), the supernatant was collected and analyzed by LC-MS/MS with the gradient of $H_2O$-0.1% formic acid/AcCN-$H_2O$ (8:2)-0.1% formic acid = 95/5 to 0/100 over 3.5 min. MMAE was quantified by multiple reaction monitoring (MRM) mode; $m/z$ = 718.5 > 134.0, 152.0, 686.4 (cone voltage = 40 V, collision energy = 30 V). For analysis of peptide products from ADC linkers, ADC linkers (30 µM) were mixed with mouse plasma (1/100 dilution) in phosphate buffer (100 mM, pH 7.4) containing CHAPS (0.1%) and incubated at 37°C for 9 h. The reaction was mixed with an equal volume of AcCN-10% formic acid. After centrifugation (14,000 rpm × 10 min, 4°C), the supernatant was collected and analyzed by LC-MS/MS with the gradient of $H_2O$-0.1% formic acid/AcCN-$H_2O$ (8:2)-0.1% formic acid = 95/5 to 0/100 over 3.5 min. The mass analysis was performed with scan mode ($ESI^+$, $m/z$ = 250-1250).

**Antibody preparation**

Antibody preparation was performed following the literature procedure[4] with modifications. Trastuzumab

(Herceptin) was dissolved in $H_2O$ to prepare 21 mg/mL solution and was diluted to 1 mg/mL (3.4 μM) in 500 μL sodium borate buffer (100 mM, pH 8.0) containing CHAPS (0.1%). 1.7 μL of Azide-PEG4-SE (TCI) solution in DMSO (10 mM) was added (final 34 μM, 10×) and mixed immediately. The reaction was incubated at 25°C for 1 h. The solution was loaded onto PD MiniTrap G-25 (Cytiva) and eluted with 1 mL Tris-HCl buffer (100 mM, pH 7.4). A 60 μL aliquot of 0.5 mg/mL (1.7 μM) solution was mixed with ADC linkers (6.8 μM; 4×), $CuSO_4$ (1 mM), BTTP (3 mM), and sodium ascorbate (3 mM) and incubated at 25°C for 3 h. The solution was diluted to 500 μL in phosphate buffered saline (PBS, pH 7.4), loaded onto PD MiniTrap G-25 (Cytiva) and eluted with 1 mL PBS. The modified antibody solution was used immediately after preparation.

**Table S1** List of sHMRG-based fluorogenic probes used in screening

| # | Sequence | Potential target | Preparation (※1) | References (※2) |
|---|---|---|---|---|
| 1 | Suc-LRR | Proteasome trypsin-like | SCCR | *ChemMedChem* **2010**, *5*, 1236-1241 |
| 2 | Suc-RR | Cathepsin B | SCCR | *Biochemistry* **2022**, *61*, 1904-1914 |
| 3 | Suc-PFR | Kallikrein | RP-MPLC | *Rinsho Kagaku* **1981**, *19*, 140-148 |
| 4 | Suc-GFR | Plasma kallikreins | SCCR | *J. Biochem.* **1977**, *82*, 1495-1498 |
| 5 | Suc-GPFR | Pancreatic kallikreins, Plasmin | SCCR | *J. Biochem.* **1977**, *82*, 1495-1498 |
| 6 | Suc-FVR | Thrombin | SCCR | *Chem. Bioll. Drug Des.* **2006**, *68*, 11-19 |
| 7 | Suc-IEGR | Factor Xa | SCCR | *J. Biochem.* **1977**, *82*, 1495-1498 |
| 8 | Suc-SGR | Factor Xa | SCCR | *J. Biochem.* **1977**, *82*, 1495-1498 |
| 9 | Suc-GGR | Urokinase | SCCR | *J. Biochem.* **1977**, *82*, 1495-1498 |
| 10 | Suc-EGR | Urokinase | SCCR | *J. Biochem.* **1977**, *82*, 1495-1498 |
| 11 | Suc-DPR | Tryptase | SCCR | *Bioorg. Med. Chem.* **2002**, *10*, 3637-3647 |
| 12 | Suc-AKR | Neurohypophysial granule endopeptidase | SCCR | *Biochem. Biophys. Res. Commun.* **1992**, *183*, 128-137 |
| 13 | Suc-IGGR | | SCCR | |
| 14 | Suc-AKK | | SCCR | |
| 15 | Suc-DPK | | SCCR | |
| 16 | Suc-FVK | | SCCR | |
| 17 | Suc-GPK | Plasmin | SCCR | *J. Biochem.* **1980**, *88*, 183-190 |
| 18 | Suc-RSLK | Site 1 Protease | RP-MPLC | *J. Biol. Chem.* **1999**, *274*, 22805-22812 |
| 19 | Suc-VLK | Calpain, Regmain | SCCR | *J. Biol. Chem.* **1984**, *259*, 12489-12494 |
| 20 | Suc-LRK | | SCCR | |
| 21 | Suc-PFK | | SCCR | |
| 22 | Suc-RTKK | | SCCR | |
| 23 | Suc-IGGK | | SCCR | |
| 24 | Suc-GGK | | SCCR | |
| 25 | Suc-EGK | | SCCR | |
| 26 | Suc-VPK | | SCCR | |
| 27 | Suc-IEGK | | SCCR | |
| 28 | Suc-SGK | | SCCR | |
| 29 | Suc-GFK | | SCCR | |
| 30 | Suc-GPFK | | SCCR | |
| 31 | Suc-RK | | SCCR | |
| 32 | Suc-FK | | SCCR | |
| 33 | Suc-VVK | | SCCR | |
| 34 | Suc-AAK | | SCCR | |
| 35 | Suc-AAVY | Proteasome chymotrypsin-like | SCCR | *ChemMedChem* **2010**, *5*, 1236-1241 |
| 36 | Suc-KGISSEY | Kallikrein 2 | SCCR | *Biochem. Biophys. Res. Commun.* **1997**, *238*, 549-555 |
| 37 | Suc-RPY | Kallikrein 3 (PSA) | SCCR | *J. Biol. Chem.* **1997**, *272*, 21582-21588 |
| 38 | Suc-KHLY | Kallikrein 7 | SCCR | *J. Invest. Dermatol.* **2017**, *137*, 430-439 |
| 39 | Suc-LLVY | Chymotrypsin, Ingensin, Proteasome, Calpain | SCCR | *ChemMedChem* **2010**, *5*, 1236-1241 |
| 40 | Suc-KPY | | SCCR | |
| 41 | Suc-KAY | | SCCR | |
| 42 | Suc-NLY | | SCCR | |
| 43 | Suc-LY | | SCCR | |
| 44 | Suc-AAY | | RP-MPLC | |
| 45 | Suc-WEHD | Caspase-1 | SCCR | *J. Biol. Chem.* **1997**, *272*, 9677-9682 |
| 46 | Suc-GYVAD | Caspase-1 | SCCR | *J. Biol. Chem.* **1997**, *272*, 9677-9682 |
| 47 | Suc-GWEHD | Caspase-1 | SCCR | *J. Biol. Chem.* **1997**, *272*, 9677-9682 |
| 48 | Suc-YVAD | Caspase-1 | SCCR | *Chem. Pharm. Bull.* **1995**, *43*, 1336-1339 |
| 49 | Suc-DYEVD | Caspase-1, Caspase-3, Caspase- 7 | SCCR | *J. Biol. Chem.* **1997**, *272*, 9677-9682 |
| 50 | Suc-GLEVD | Caspase-1, Caspase- 6 | SCCR | *J. Biol. Chem.* **1997**, *272*, 9677-9682 |
| 51 | Suc-GDEVD | Caspase-1, Caspase- 3, Caspase-7 | SCCR | *J. Biol. Chem.* **1997**, *272*, 9677-9682 |
| 52 | Suc-VVEID | Caspase-1, Caspase-3, Caspase-6, Caspase-7 | SCCR | *J. Biol. Chem.* **1997**, *272*, 9677-9682 |
| 53 | Suc-VDQQD | Caspase-2 | SCCR | *J. Biol. Chem.* **1997**, *272*, 9677-9682 |
| 54 | Suc-VDVAD | Caspase-2, Caspase-3 | SCCR | *J. Biol. Chem.* **1997**, *272*, 9677-9682 |
| 55 | Suc-DESD | Caspase-2, Caspase-3 | SCCR | *J. Biol. Chem.* **1997**, *272*, 9677-9682 |
| 56 | Suc-DEVD | Caspase-3 | RP-MPLC | *J. Biol. Chem.* **1997**, *272*, 9677-9682 |
| 57 | Suc-GVQVD | Caspase-3, Caspase-6 | SCCR | *J. Biol. Chem.* **1997**, *272*, 9677-9682 |
| 58 | Suc-IETD | Caspase-8, Granzyme B | SCCR | *Cell Death Differ.* **2008**, 15, 322-331 |
| 59 | Suc-GIETD | Caspase-8, Granzyme B | SCCR | *Cell Death Differ.* **2008**, 15, 322-331 |
| 60 | Suc-LEHD | Caspase-9 | SCCR | *Br. J. Ophthalmol.* **2006**, *90*, 760-764 |
| 61 | Suc-LLEHD | Caspase-9 | SCCR | *Br. J. Ophthalmol.* **2006**, *90*, 760-764 |
| 62 | Suc-LEED | Caspase-13 | SCCR | *Blood* **2004**, *104*, 1998 |
| 63 | Suc-IEPD | Granzyme B | SCCR | *J. Am. Chem. Soc.* **2020**, *142*, 7075-7082 |
| 64 | Suc-GIEPD | Granzyme B | SCCR | *J. Am. Chem. Soc.* **2020**, *142*, 7075-7082 |
| 65 | Suc-EETD | | SCCR | |
| 66 | Suc-VETD | | RP-MPLC | |
| 67 | Suc-IKTD | | SCCR | |
| 68 | Suc-ISTD | | SCCR | |
| 69 | Suc-IEKD | | SCCR | |
| 70 | Suc-IEVD | | SCCR | |
| 71 | Suc-ELEED | | SCCR | |
| 72 | Suc-EGITD | | SCCR | |
| 73 | Suc-VWTTD | | RP-MPLC | |
| 74 | Suc-AAPPD | | SCCR | |
| 75 | Suc-AAD | | SCCR | |
| 76 | Suc-LLE | Proteasome | SCCR | *ChemMedChem* **2010**, *5*, 1236-1241 |
| 77 | Suc-IETE | | SCCR | |
| 78 | Suc-GDEVE | | SCCR | |
| 79 | Suc-DYEVE | | SCCR | |
| 80 | Suc-GIETE | | SCCR | |
| 81 | Ac-GP | FAP-alpha | SCCR | *bioRxiv* (DOI: 10.1101/2025.08.07.669035) |
| 82 | Suc-GPLGP | Amyloid A4 generating enzyme | SCCR | *FEBS Lett.* **1990**, *260*, 131-134 |
| 83 | Suc-AAA | Elastase | SCCR | *Cell. Rep. Methods* **2024**, *4*, 100688 |
| 84 | Suc-VIA | | SCCR | |
| 85 | Suc-AAPAbu | Pancreatic elastase, neutrophil elastase | SCCR | *Cell. Rep. Methods* **2024**, *4*, 100688 |
| 86 | Suc-AAPV | Elastase | SCCR | *J. Pathol. Microbiol. Immun.* **1992**, *100*, 1073-1080 |
| 87 | Suc-AAV | | SCCR | |
| 88 | Suc-AAI | | SCCR | |
| 89 | Suc-AAH | | SCCR | |
| 90 | Suc-AAS | | SCCR | |
| 91 | Suc-EVCit | Cathepsins | RP-MPLC | *Nat. Commun.* **2018**, *9*, 2512 |
| 92 | Suc-LRGG | SUMO-specific protease | SCCR | *Cell. Rep. Methods* **2024**, *4*, 100688 |
| 93 | Suc-QTGG | SUMO-specific protease | SCCR | *Cell. Rep. Methods* **2024**, *4*, 100688 |
| 94 | $\gamma$Glu | GGT1 | SCCR | |
| 95 | R | CD13 | SCCR | *Cell. Rep. Methods* **2024**, *4*, 100688 |
| 96 | GP | DPP4 | RP-MPLC | *Cell. Rep. Methods* **2024**, *4*, 100688 |
| 97 | SP | DPP4, DPP9 | SCCR | *RSC Chem. Biol.* **2022**, *3*, 859-867 |
| 98 | HA | | SCCR | |
| 99 | KA | | SCCR | |
| 100 | GA | | SCCR | |
| 101 | YA | | SCCR | |
| 102 | Ac-A | APEH | SCCR | *J. Am. Chem. Soc.* **2013**, *135*, 6002-6005 |
| 103 | Ac-M | APEH | SCCR | *J. Am. Chem. Soc.* **2013**, *135*, 6002-6005 |

※1: Preparation refers to the final process employed to acquire the compounds used in the study. SCCR means that the probes were used after the release from the solid phase without further purification. RP-MPLC means that the probes were purified over reverse-phase MPLC after the release from the solid phase.
※2: References were primarily derived from literature discussing the substrate specificities of 7-amino-4-methylcoumarin (AMC) or *p*-nitrophenylaniline (*p*-NA).

**Table S2. Category of proteases and representative enzymes**

| Category | P1 amino acids | Representative enzymes |
| --- | --- | --- |
| Trypsin-like | Arg, Lys | Kallikreins, Cathepsins, Trypsin, Plasmin, Thrombin, Factor Xa, Furin, Urokinase, Enterokinase, Proteasome (trypsin-like), Tryptase, Calpain |
| Chymotrypsin-like | Phe, Trp, Tyr | Kallikreins, Cathepsins, Chymotrypsin, Chymase, Calpain, Renin, Proteasome (chymotrypsin-like) |
| Caspase-like | Asp | Caspase-1, Caspases, Granzyme B, Proteasome (caspase-like) |
| Elastase-like | Ala, Leu, Val | Neutrophil elastase, Pancreatic elastase |
| Pro endopeptidase | Pro | DPPs, FAP$\alpha$, Amyloid A4 generating enzyme |
| SENPs | Gly | SENP1, SENP2 |

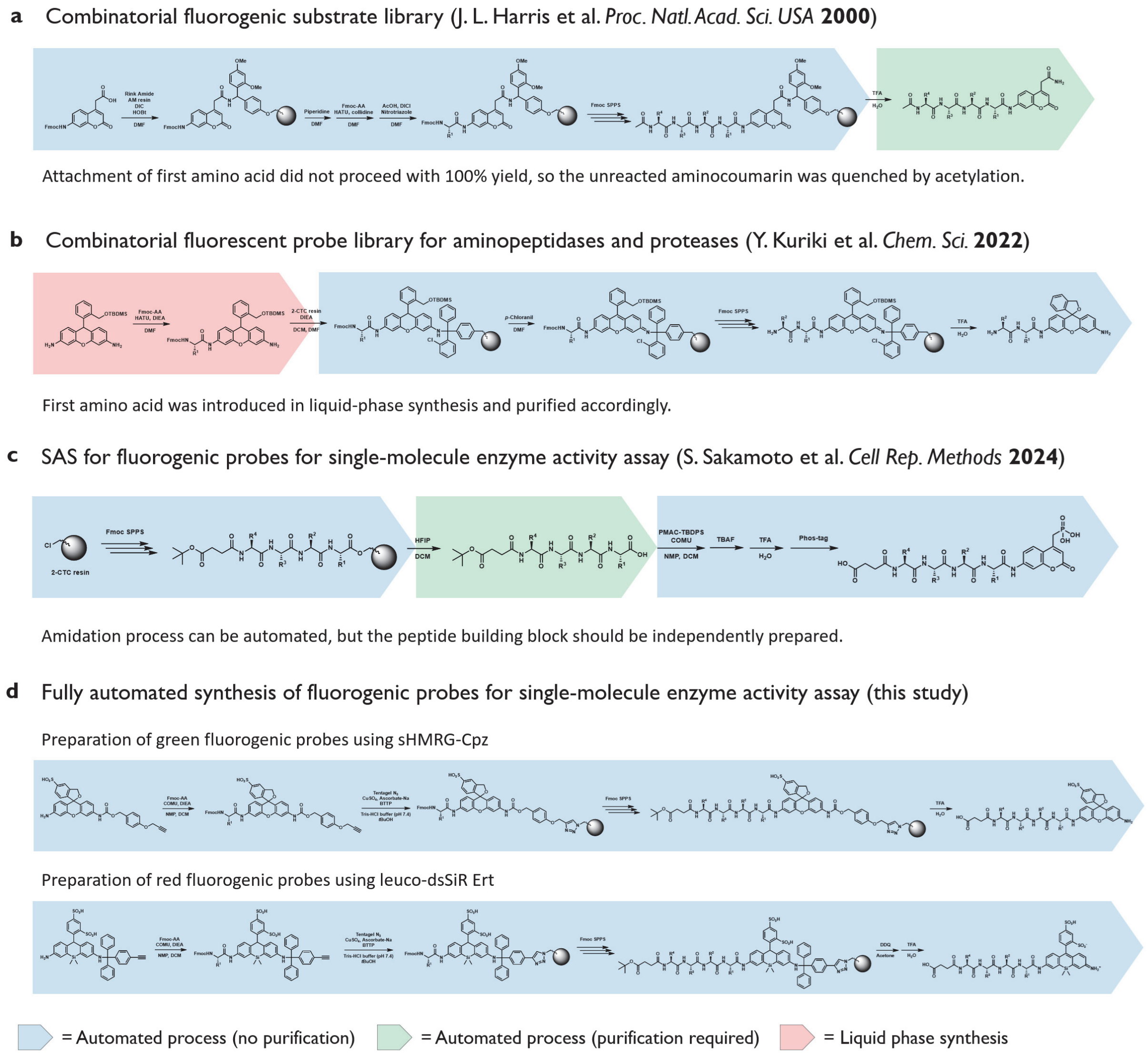


**Figure S1.** Comparison of methodologies used to prepare the library of fluorogenic substrates for peptidases. (a)–(c) refer to previously reported procedures for the preparation of peptide-modified fluorogenic probes for peptidases[5–7]. (c) represents the SAS strategy, and (d) represents the SCCR strategy employed in this study. Blue highlights indicate steps that can be automated using a peptide synthesizer; yellow highlights indicate steps that can be performed in an automated manner but require manual purification to achieve assay-ready purity (>90%); and red highlights indicate steps performed using standard organic synthesis procedures.

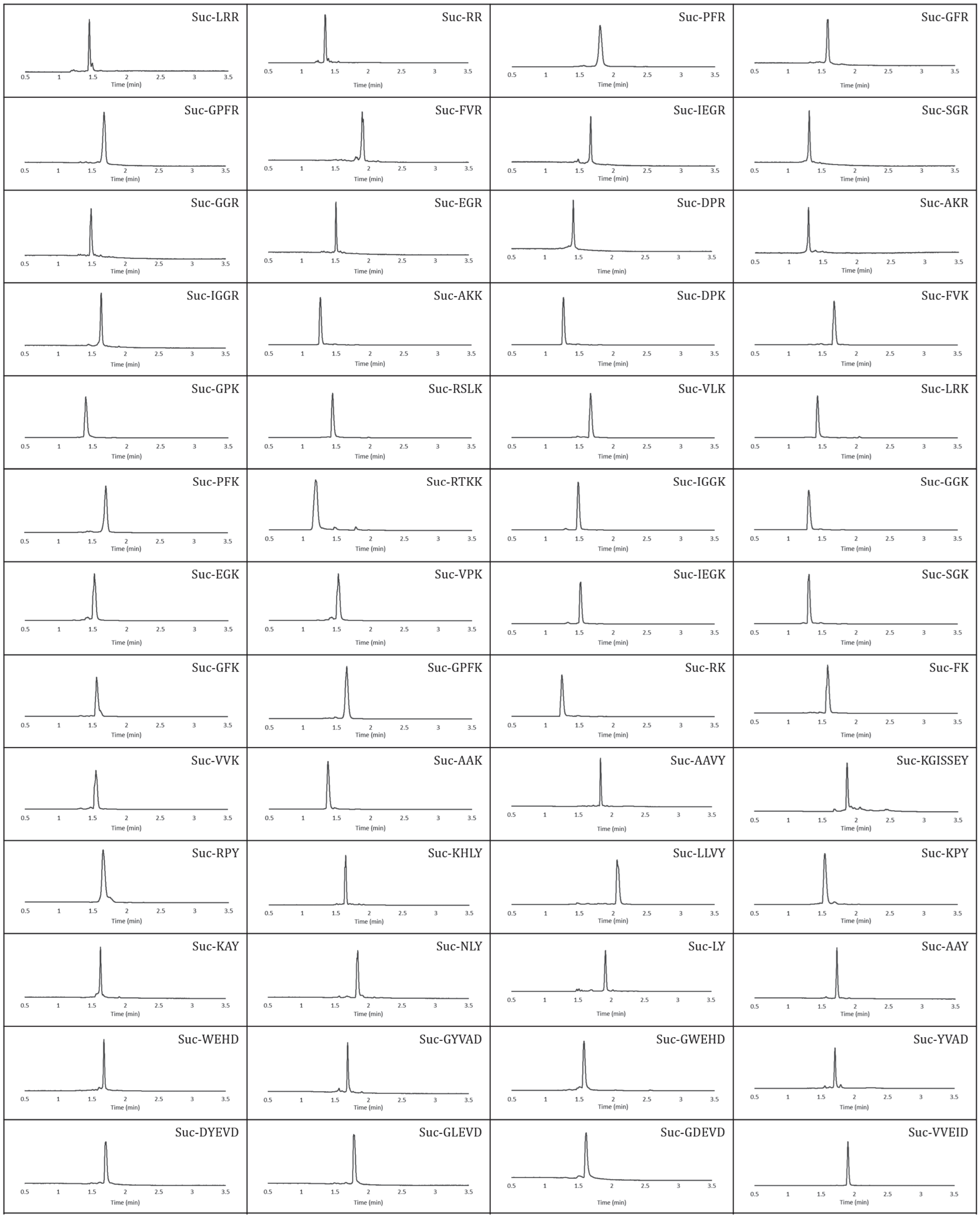


**Figure S2.** LC chromatograms (absorbance at 500 nm) of sHMRG-based probes #1-#53. Chromatograms for previously reported compounds[8] were not included.

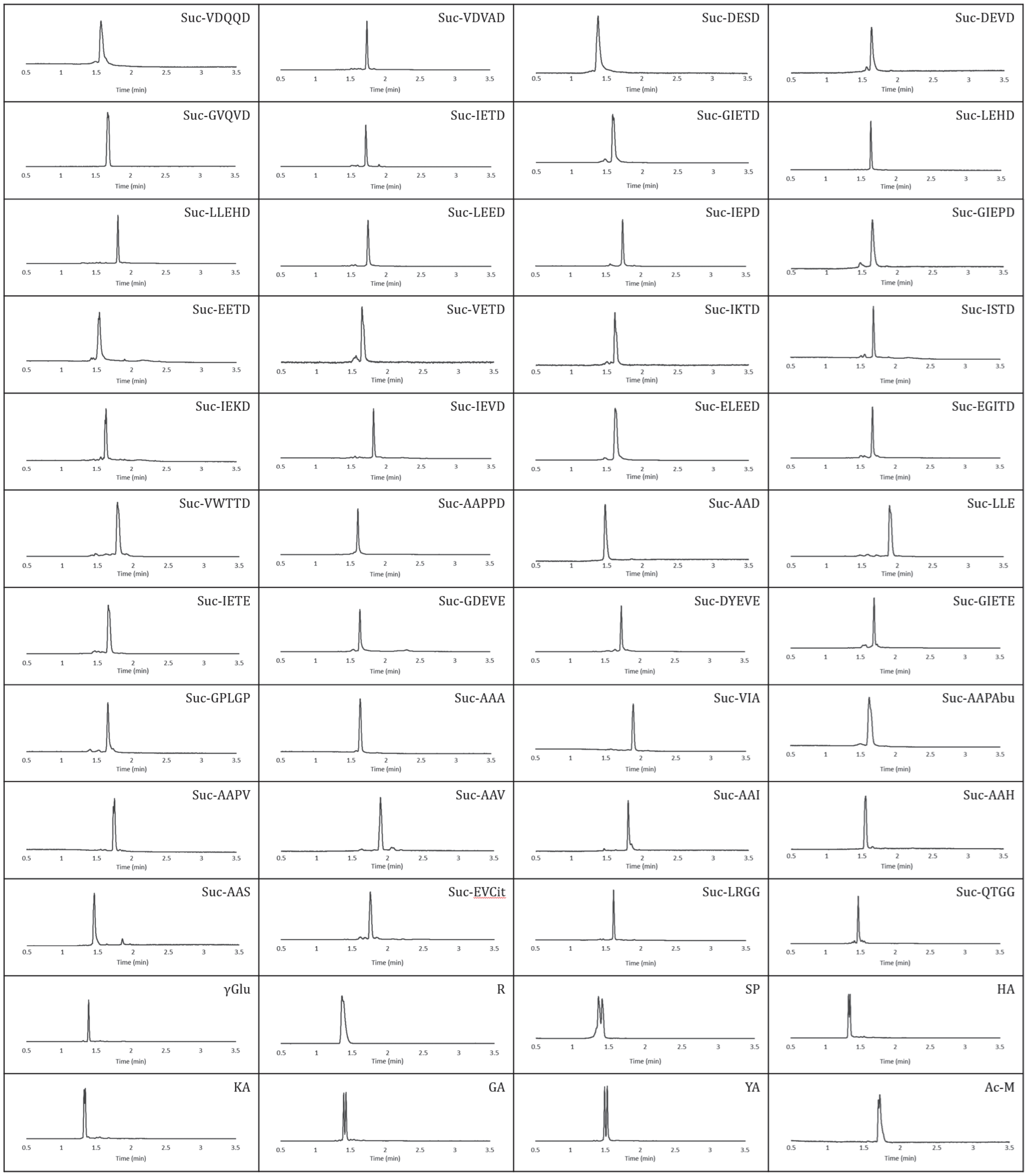


**Figure S3.** LC chromatograms (absorbance at 500 nm) of sHMRG-based probes #54-#103. Chromatograms for previously reported compounds[8] were not included.

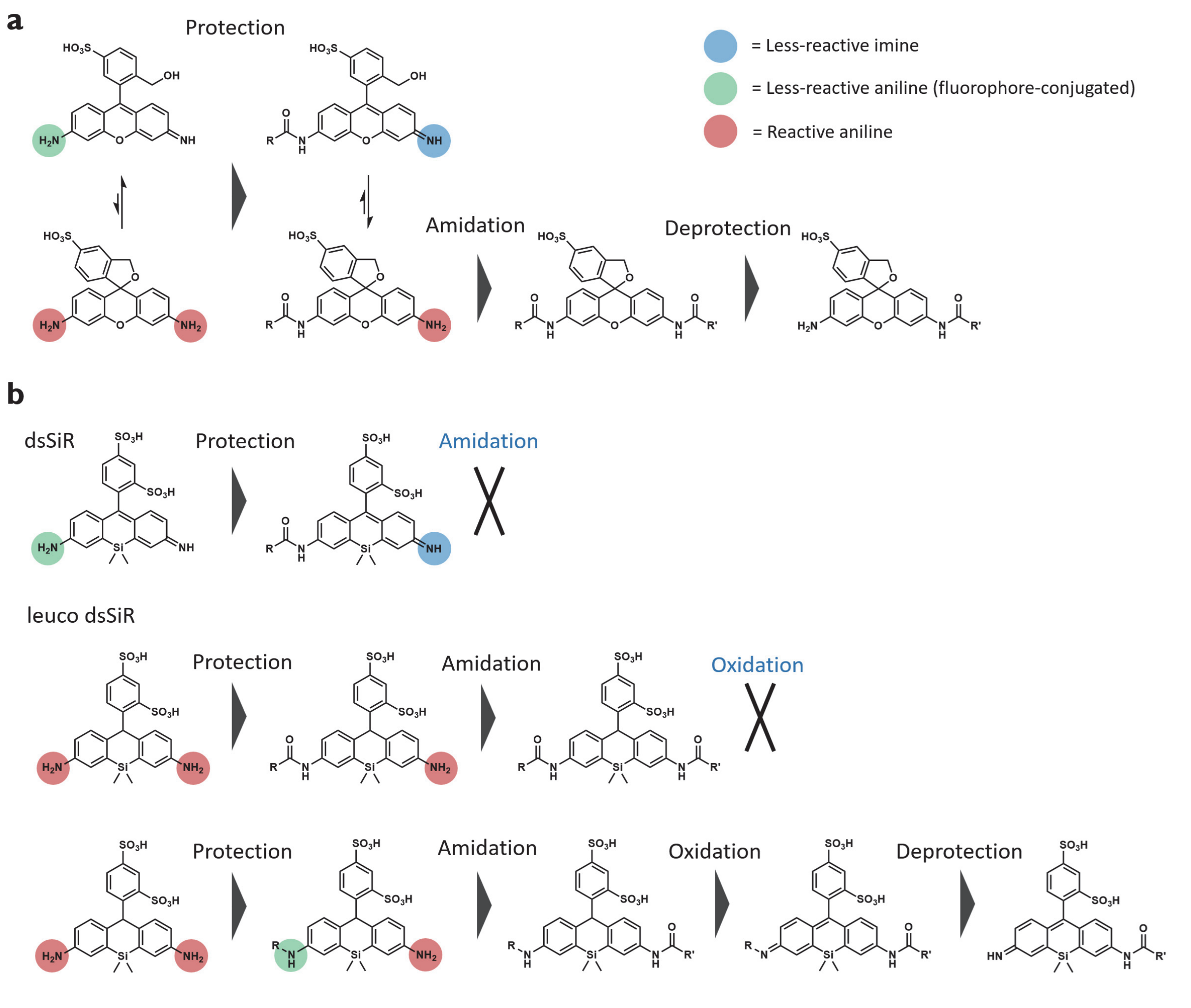


**Figure S4.** Requirements for appropriate protecting groups to enable reactions at the aniline moieties of rhodamine fluorophores. (a) Protection strategy of sHMRG. Red circles indicate anilines with high reactivity (not part of the extended conjugated system of the fluorophore), yellow circles indicate anilines with low reactivity (within the conjugated system of the fluorophore), and blue circles indicate imines that are unreactive toward amidation. (b) Protection strategy of dsSiR and its leuco form for the design of SCCR-compatible starting materials.

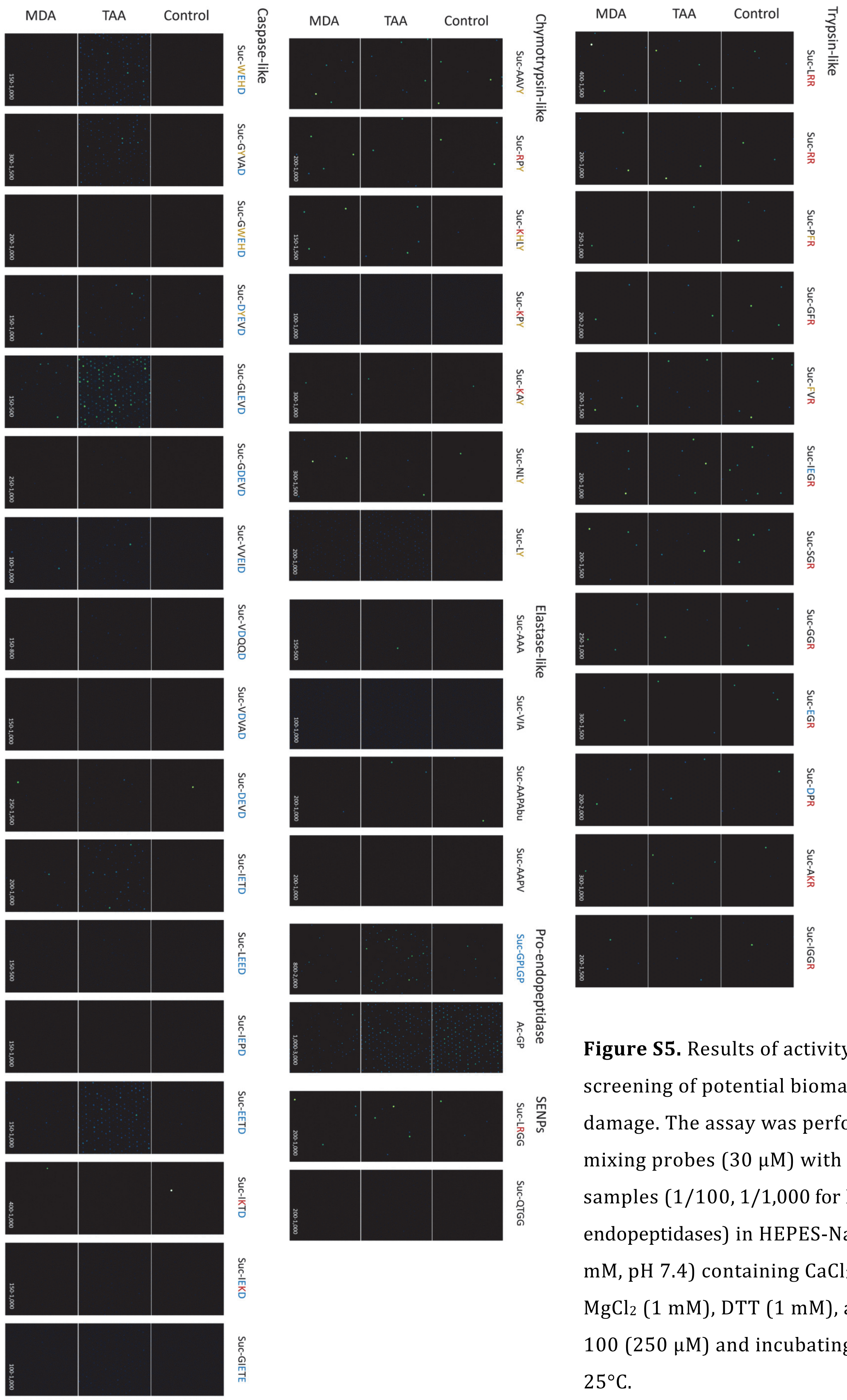


**Figure S5.** Results of activity-based screening of potential biomarkers of liver damage. The assay was performed by mixing probes (30 μM) with plasma samples (1/100, 1/1,000 for Pro-endopeptidases) in HEPES-Na buffer (100 mM, pH 7.4) containing $CaCl_2$ (1 mM), $MgCl_2$ (1 mM), DTT (1 mM), and Triton X-100 (250 μM) and incubating for 18 h at 25°C.

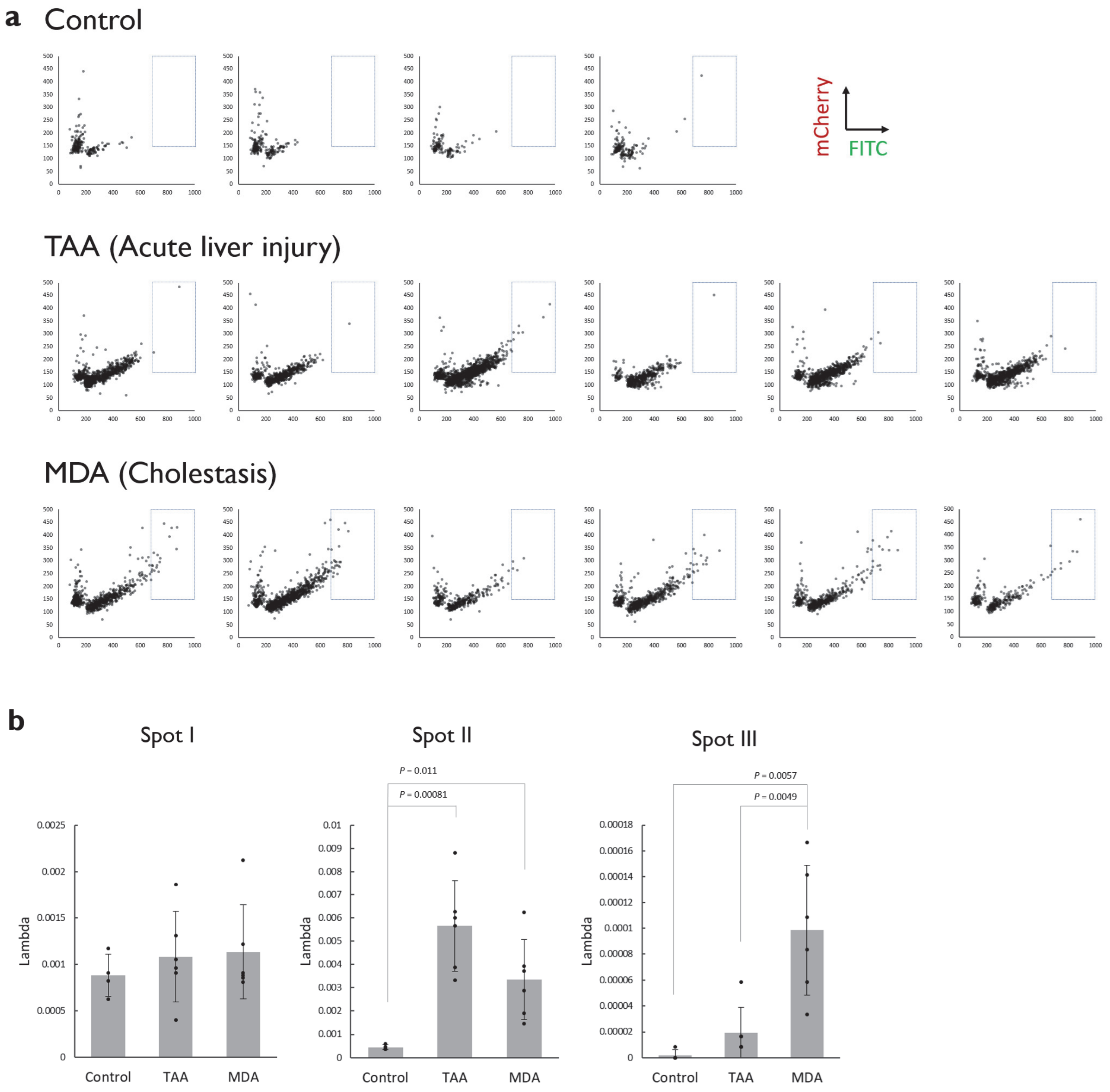


**Figure S6.** Analysis of individual blood samples by Suc-LHLY-sHMRG and Suc-AAVY-dsSiR. The same conditions as those in **Figure 2f** and **2g** were applied. (a) Scatter plot of the fluorescence intensities of wells containing enzymes. Spot III is indicated as a blue rectangle. (b) Dot plot of the number of enzyme species categorized into each spot. The thresholds for the spots are shown in **Figure 2g**. n = 4 for control and n = 6 for TAA- or MDA-treated mice. Error bars represent S. D.

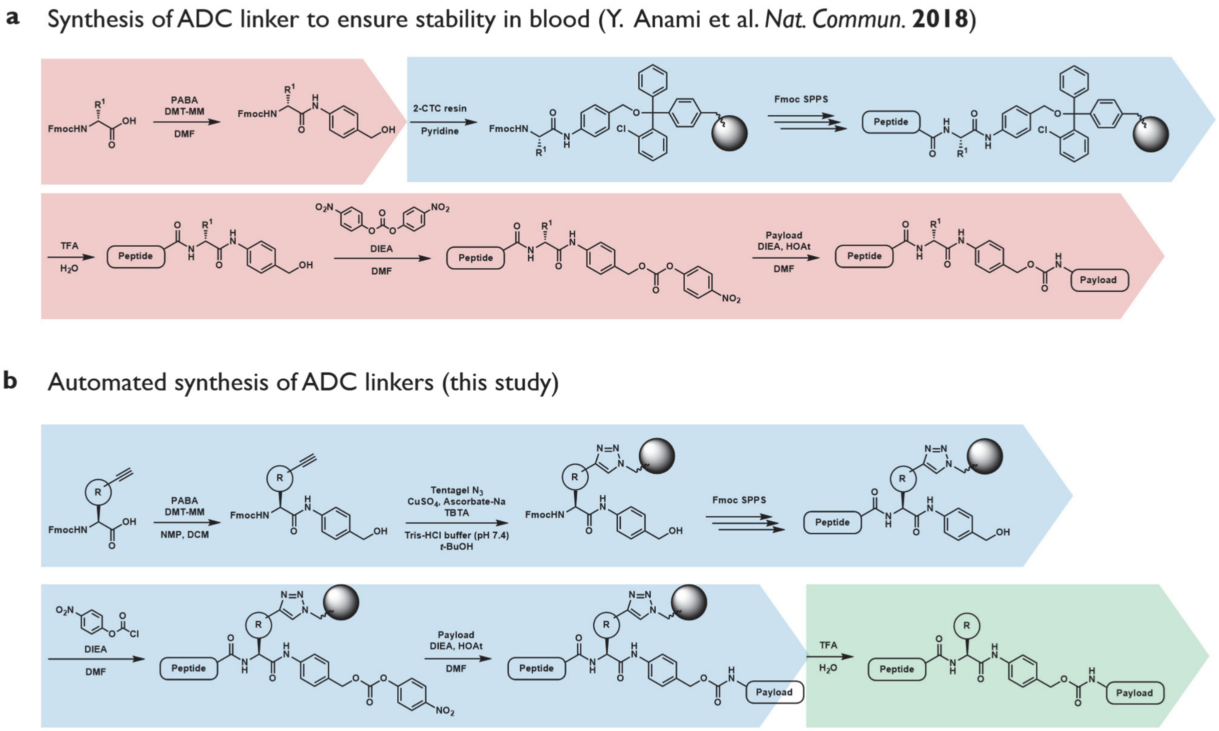


**Figure S7.** Comparison of methodologies for preparing ADC linkers. (a) refers to the reported procedures for the conventional scheme of ADC linker preparation[9]. (b) refers to the SCCR strategy employed in this study.

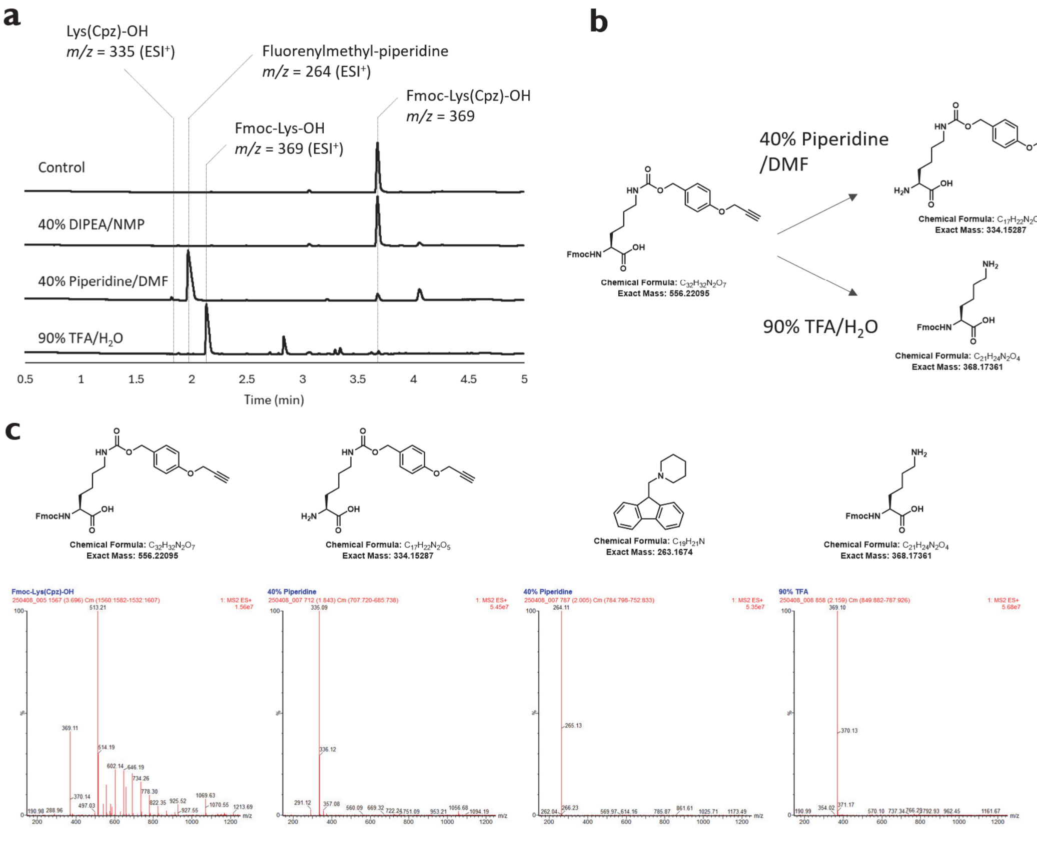


**Figure S8.** Reactivity of Fmoc-Lys(Cpz)-OH under various reaction conditions. (a) Chromatograms (254 nm) of Fmoc-Lys(Cpz)-OH (100 μM) after incubation at 25°C for 1 h in various conditions used in peptide synthesis. The representative mass values observed for each peak are shown. (b) Representative deprotection routes of Fmoc-Lys(Cpz)-OH. (c) Mass spectra of peaks observed in (a).

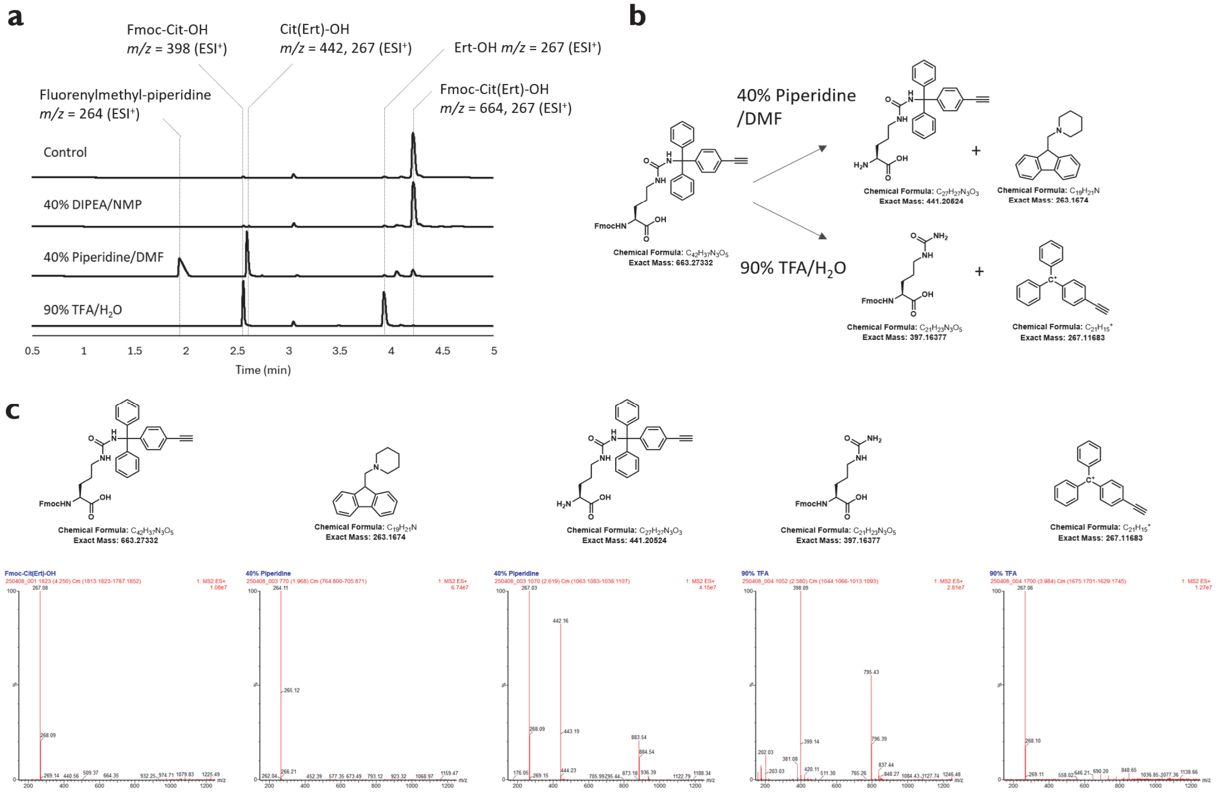


**Figure S9.** Reactivity of Fmoc-Cit(Ert)-OH under various reaction conditions. (a) Chromatograms (254 nm) of Fmoc-Cit(Ert)-OH (100 μM) after incubation at 25°C for 1 h in various conditions used in peptide synthesis. The representative mass values observed for each peak are shown. (b) Representative deprotection routes of Fmoc-Cit(Ert)-OH. (c) Mass spectra of peaks observed in (a).

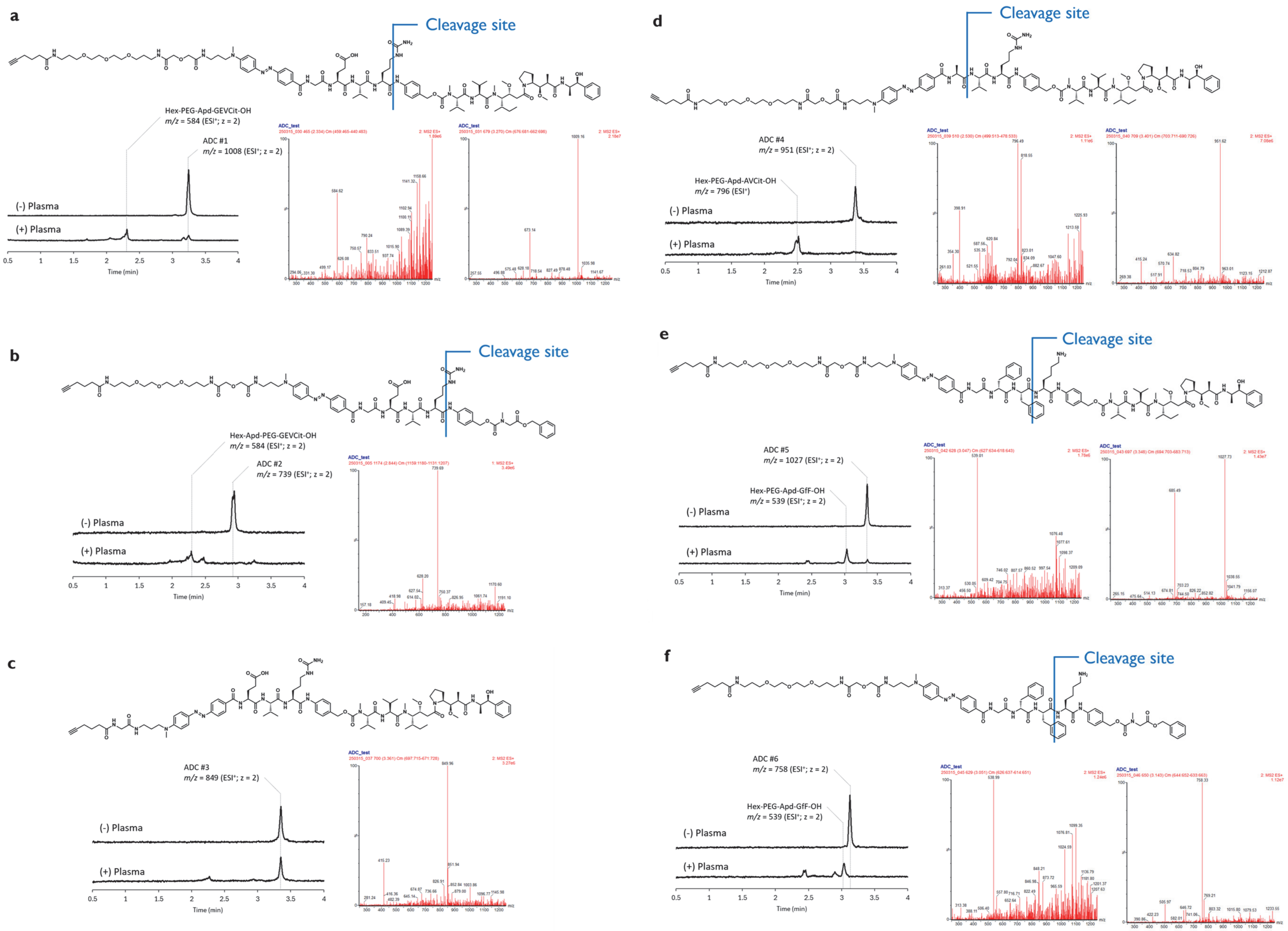
a
Cleavage site
Hex-PEG-Apd-GEVCit-OH
m/z = 584 (ESI+; z = 2)
ADC #1
m/z = 1008 (ESI+; z = 2)
(-) Plasma
(+) Plasma
Time (min)
b
Cleavage site
Hex-Apd-PEG-GEVCit-OH
m/z = 584 (ESI+; z = 2)
ADC #2
m/z = 739 (ESI+; z = 2)
(-) Plasma
(+) Plasma
Time (min)
c
ADC #3
m/z = 849 (ESI+; z = 2)
(-) Plasma
(+) Plasma
Time (min)
d
Cleavage site
ADC #4
m/z = 951 (ESI+; z = 2)
Hex-PEG-Apd-AVCit-OH
m/z = 796 (ESI+)
(-) Plasma
(+) Plasma
Time (min)
e
Cleavage site
ADC #5
m/z = 1027 (ESI+; z = 2)
Hex-PEG-Apd-GfF-OH
m/z = 539 (ESI+; z = 2)
(-) Plasma
(+) Plasma
Time (min)
f
Cleavage site
ADC #6
m/z = 758 (ESI+; z = 2)
Hex-PEG-Apd-GfF-OH
m/z = 539 (ESI+; z = 2)
(-) Plasma
(+) Plasma
Time (min)

**Figure S10.** Cleavage of ADC linkers in mouse plasma. The reaction was performed by mixing ADC linkers (30 μM) with mouse plasma (1/100) in phosphate buffer (100 mM, pH 7.4) and incubating at 37°C for 9 h. The chromatograms (500 nm) show the results with or without plasma samples. The results for (a) ADC linker #1, (b) ADC linker #2, (c) ADC linker #3, (d) ADC linker #4, (e) ADC linker #5, and (f) ADC linker #6 are shown.

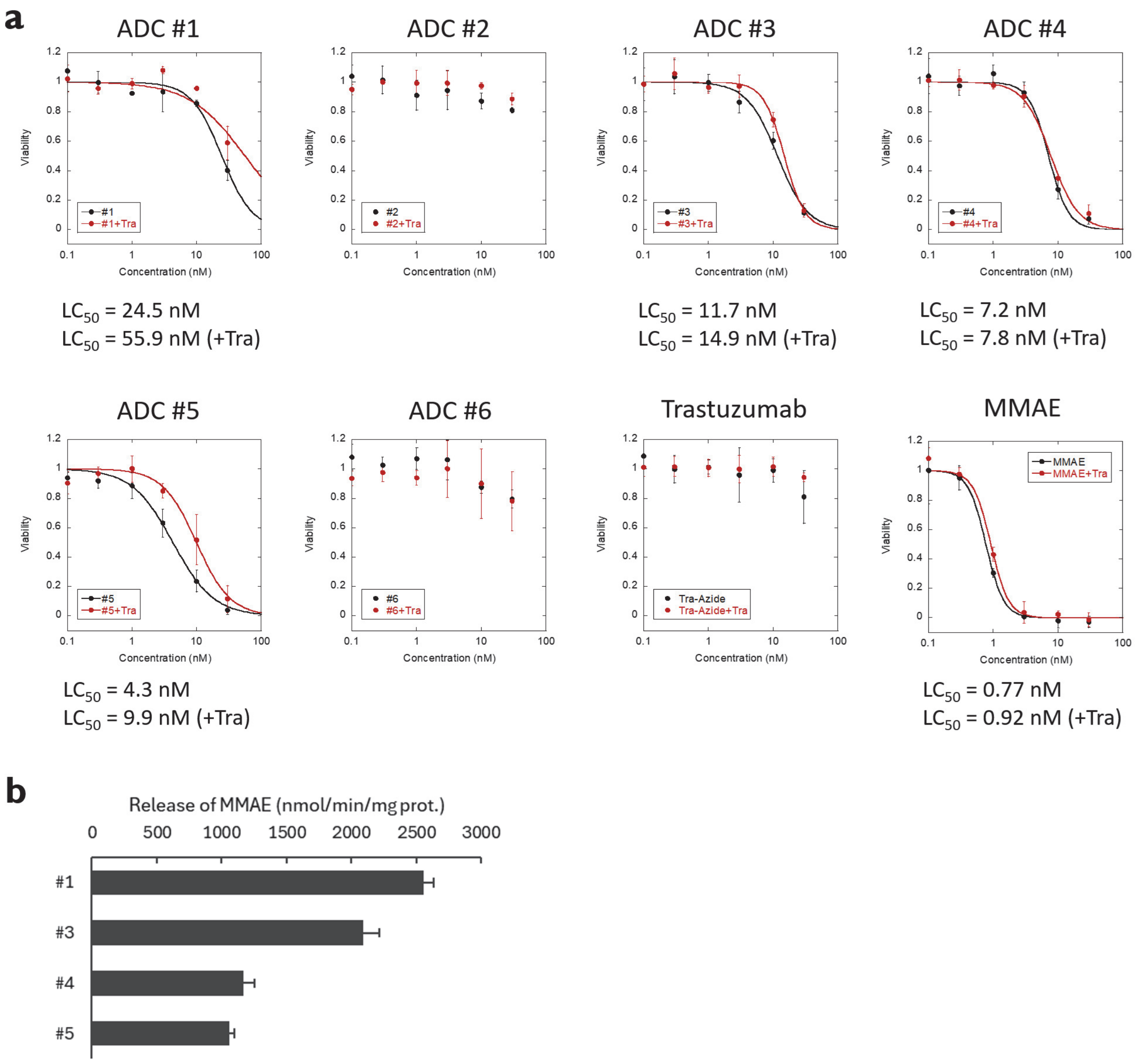


**Figure S11.** Activity of ADCs in SKBR3 breast tumor cells. (a) $LC_{50}$ values of trastuzumab conjugated with ADC linkers toward SKBR3 cells. In competing conditions (+Tra, red lines), the ADC was co-incubated with trastuzumab (100 nM). Error bars represent S. D. (n = 3). (b) Formation of MMAE after incubating ADC linkers (#1, #3, #4, and #5, 30 μM) with SKBR3 cell lysate (0.15 mg/mL) incubated in phosphate buffer (100 mM, pH 5.5) containing DTT (1 mM) and CHAPS (0.1%) for 20 h. Error bars represent SD (n = 3).

#### Supplementary methods for synthesis and characterization of compounds

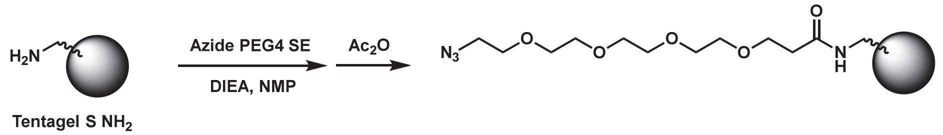


**Scheme S1.** Preparation of Tentagel-azide

#### Preparation of Tentagel-azide

Tentagel S $NH_2$ (Sigma-Aldrich 86364, 90 μm, 0.26 mmol/g, 600 mg, 156 μmol) was suspended in a mixture of DIEA and NMP (5 mL, 1:2, v/v), followed by the addition of Azide-$PEG_4$-SE (TCI A2388, 25 mg, 64 μmol). The mixture was stirred at 25°C for 18 h. After the reaction, the beads were washed three times with DMF, then resuspended in a mixture of DIEA and NMP (5 mL, 1:2, v/v), and acetic anhydride (156 μL) was added. The mixture was stirred at 25°C for 3 h. The beads were then washed three times with DMF and three times with DCM, dried under reduced pressure, and stored at 4°C.

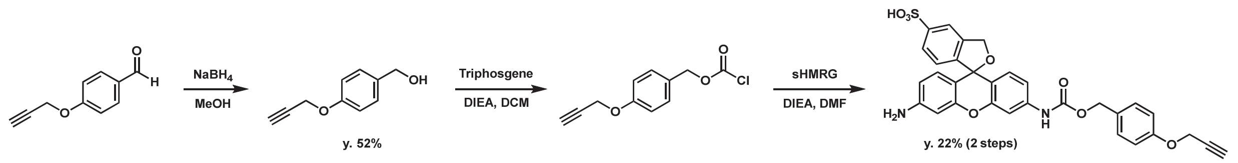


**Scheme S1.** Preparation of Cpz-OH and protection of sHMRG.

#### Synthesis of *p*-Propargyloxy benzyl alcohol (Pzl-OH, 1)

*p*-Propargyloxy benzaldehyde (2.5 g, 15.6 mmol) was dissolved in methanol (10 mL) and stirred on ice. Sodium borohydride ($NaBH_4$, 1.18 g, 31 mmol, 2 eq.) was added portionwise, and the mixture was stirred at 25°C for 30 min. After the reaction, water (20 mL) was added, and the product was extracted three times with dichloromethane (DCM), then washed once with brine. The organic layer was dried over $Na_2SO_4$ and evaporated to afford a colorless liquid (1.76 g, yield: 69%). The identity of the product was confirmed by comparison of the LC retention time and MS signal with those of an independently synthesized compound via propargylation of *p*-hydroxybenzyl alcohol, as reported[10].

LRMS ($ESI^+$): $m/z$ = 145 ($[M–OH]^+$)

#### Synthesis of sHMRG-Cpz (2)

sHMRG is prepared according to the literature[8]. Pzl-OH (150 mg, 0.92 mmol) was dissolved in THF (5 mL), and DIEA (805 μL, 4.62 mmol, 5 equiv.) was added under an Ar atmosphere at 0°C. A solution of triphosgene (69 mg, 0.23 mmol, 0.25 equiv.) in THF (1 mL) was slowly added. The mixture was stirred at 25°C for 30 min to afford a chloroformate solution (A). sHMRG (20 mg, 0.05 mmol) was dissolved in DMSO (1 mL) and DIEA (300 μL), and solution (A) was added in 500 μL portions. The reaction was monitored by HPLC every 10 min, and upon ~50% conversion of sHMRG to the Cpz-protected derivative, the mixture was quenched with water

(1 mL) and concentrated under reduced pressure. The residue was dissolved in water and purified by preparative MPLC (eluent: A = 0.1% TFA in $H_2O$, B = 0.1% TFA in MeCN, gradient from A:B = 95:5 to 5:95 over 15 min). The desired fractions were pooled and lyophilized to give an orange solid (7 mg, yield: 24%).

$^1$H-NMR (400 MHz, $CD_3OD$): δ 3.45 (s, 1H), 4.33 (s, 2H), 4.71 (s, 2H), 5.18 (s, 2H), 6.7–7.1 (m, 6H), 7.2–7.4 (m, 5H), 7.95 (d, 1H, $J$ = 8.2 Hz), 8.16 (s, 1H).

$^{13}$C NMR (100 MHz, DMSO-$d_6$): δ55.9, 61.3, 67.2, 78.9, 79.7, 97.7, 104.3, 115.0, 115.4, 117.8, 124.9, 125.7, 128.4, 129.0, 129.2, 130.1, 130.8, 131.1, 133.9, 135.8, 140.8, 150.3, 153.6, 155.6, 157.8, 158.5, 159.7, 162.9.

HRMS (ESI$^+$): $m/z$ Calcd. for $[M+H]^+$, 585.1326, Found, 585.1351 (+2.5 mDa).

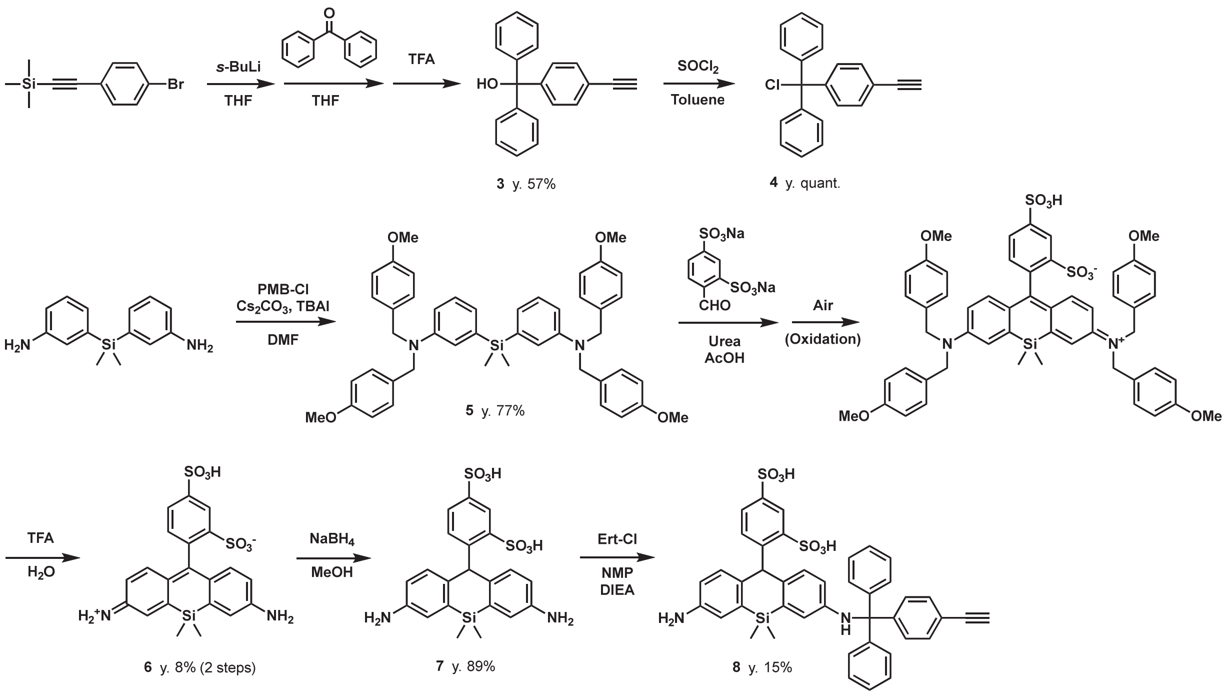


**Scheme S2.** Preparation of Ert-OH and leuco dsSiR-Ert.

**Synthesis of Ethynyl-trityl alcohol (Ert-OH, 3)**

(4-Bromophenylethynyl)-trimethylsilane (500 mg, 1.97 mmol) was dissolved in THF (50 mL) and cooled to -78°C under an Ar atmosphere. *sec*-BuLi in hexane (1.3 M, 1.5 mL, 1.97 mmol, 1 eq.) was added dropwise. The reaction mixture was allowed to warm to 25 °C and stirred for 30 min, then cooled again to -78 °C. A solution of benzophenone (360 mg, 1.97 mmol, 1 eq.) in THF (2 mL) was added dropwise, and the mixture was stirred at 25 °C for 2 h. Water was then added, and the product was extracted three times with DCM and washed once with brine. The organic layer was dried over $Na_2SO_4$ and concentrated under reduced pressure. The crude product was purified by silica gel column chromatography (eluent: hexane–DCM, 100:0 to 0:100 over 15 min) to give **3** as a colorless liquid (320 mg, yield: 57%).

$^1$H-NMR (400 MHz, $CDCl_3$): δ 2.77 (s, 1H), 3.05 (s, 1H), 7.3 (m, 12H), 7.45 (d, 2H, $J$ = 8.2 Hz)

**Synthesis of Ethynyl-trityl chloride (Ert-Cl, 4)**

Ert-OH (50 mg, 0.18 mmol) was dissolved in toluene (1 mL), and thionyl chloride ($SOCl_2$, 76 µL, 1.1 mmol, 6 eq.) was added. The reaction was refluxed for 18 h. After cooling, excess toluene was added and solvents were removed *in vacuo*. The crude product was used directly in the next step.

$^1$H-NMR (400 MHz, $CDCl_3$): δ 3.08 (s, 1H), 7.2 (m, 6H), 7.3 (m, 6H), 7.42 (d, 2H, *J* = 8.2 Hz)

**Preparation of 5**

3,3'-(Dimethylsilanediyl)dianiline was prepared according to the literature[11]. 3,3'-(Dimethylsilanediyl)dianiline (400 mg, 1.65 mmol), *p*-methoxybenzyl chloride (1.1 mL, 8.25 mmol, 5 eq.), DIEA (1.4 mL, 8.25 mmol, 5 eq.), tetrabutylammonium iodide (30 mg, 0.08 mmol, 0.05 eq.) were dissolved in DMF (2 mL) and stirred at 70°C for 6 h. The reaction was diluted by DCM and directly subjected to column for silica gel column chromatography (eluent: hexane/AcOEt) to afford compound **5** (914 mg, yield: 77%).

$^1$H-NMR (400 MHz, $CDCl_3$): δ 0.32 (s, 6H), 3.77 (s, 12H), 4.49 (s, 8H), 6.72 (dd, 2H, *J* = 8.6, 2.4 Hz), 6.76 (d, 2H, *J* = 8.4 Hz), 6.81 (d, 8H, *J* = 8.4 Hz), 6.91 (d, 2H, *J* = 2.4 Hz), 7.1-7.2 (m, 10H).

$^{13}$C-NMR (100 MHz, $CDCl_3$): δ -2.43, 53.7, 55.4, 113.8, 113.0, 114.0, 118.4, 124.4, 128.1, 128.6, 130.8, 139.0, 148.5, 158.6.

LRMS (ESI$^+$): *m/z* = 723 ([M + H]$^+$)

**Synthesis of dsSiR (6)**

Compound **5** (490 mg, 0.68 mmol), 4-formylbenzene-1,3-disulfonic acid disodium salt (142 mg, 0.81 mmol, 1.2 eq.) and urea (30 mg, 0.51 mmol, 0.75 eq.) were dissolved in acetic acid (10 mL) and stirred at 95°C for 20 h in open air. LC-MS-based reaction monitoring suggested complete consumption of the starting material, formation of the oxidated fluorophore, and partial deprotection of PMB groups. The solvents were removed *in vacuo*, and the products were purified over silica gel column chromatography (DCM/MeOH 100/0 to 20/80 over 20 min). Fractions containing tetra-PMB and tri-PMB dsSiR were collected and the solvent was removed *in vacuo*. To the remaining TFA (2 mL) was added to the residue, and the mixture waas stirred at 25°C for 40 min. LC-MS-based reaction monitoring suggested formation of dsSiR and its mono-PMB form. Solvent was removed *in vacuo*, and the products were purified over prep. MPLC (eluent: A = 0.1% triethylamine in $H_2O$, B = MeOH, gradient from A:B = 95:5 to 5:95 over 15 min) to afford dsSiR (25 mg, yield: 8%).

$^1$H NMR (400 MHz, $CD_3OD$): δ 0.47 (s, 6H), 6.54 (d, 2H, *J* = 8.4 Hz), 6.93 (d, 2H, *J* = 8.4 Hz), 7.16 (s, 2H), 7.20 (d, 1H, *J* = 7.2 Hz), 7.96 (d, 1H, *J* = 7.2 Hz), 8.44 (s, 1H).

HRMS (ESI$^+$): *m/z* Calcd. for [M+H]$^+$, 489.0605, Found, 489.0647 (+4.2 mDa).

**Synthesis of leuco-dsSiR (7)**

dsSiR (48 mg, 0.10 mmol) was dissolved in methanol (3 mL) and sodium borohydride (22 mg, 0.59 mmol) was added, and the reaction was stirred at 25°C for 2 h. $H_2O$ was added, and the solvent was removed under reduced pressure. The crude product was purified over preparative MPLC (eluent: A = 0.1% triethylamine in $H_2O$, B = MeOH, gradient from A:B = 95:5 to 5:95 over 15 min), and the combined fractions were freeze-dried to afford leuco dsSiR (43 mg, yield: 89%).

$^1$H NMR (400 MHz, $CD_3OD/D_2O$ = 1:1): δ 0.52 (s, 3H), 0.53 (s, 3H), 6.59 (dd, $J$ = 8.4, 2.4 Hz, 2H), 6.98 (d, $J$ = 8.4 Hz, 2H), 7.21 (d, $J$ = 2.4 Hz, 2H), 7.26 (d, $J$ = 7.6 Hz, 1H), 8.01 (dd, $J$ = 7.6, 2.0 Hz, 1H), 8.48 (d, $J$ = 2.0 Hz, 1H).

HRMS (ESI$^+$): $m/z$ = 491 ([M + H]$^+$)

**Synthesis of leuco dsSiR-Ert (8)**

Leuco dsSiR (20 mg, 0.04 mmol) was dissolved in NMP (20 μL). A solution of Ert-Cl (6.2 mg, 0.02 mmol, 0.5 eq.) and DIEA (36 μL, 0.2 mmol, 5 eq.) in DCM (100 μL) was added dropwise. DCM was removed under reduced pressure over 30 min. The resulting residue was dissolved in water and purified by preparative MPLC (eluent: A = 0.1% triethylamine in $H_2O$, B = MeOH, gradient from A:B = 95:5 to 5:95 over 15 min). Fractions containing the desired product were pooled. Triethylamine (50 μL) was added to the solution and lyophilized to afford leuco dsSiR Ert (4.5 mg, yield: 15%).

$^1$H NMR (400 MHz, DMSO-$d_6$): δ 0.52 (s, 6H), 3.06 (s, 1H), 4.09 (s, 1H), 6.32 (dd, 1H, $J$ = 8.0, 2.4 Hz), 6.37 (d, 1H, $J$ = 2.4 Hz), 6.48 (dd, 1H, $J$ = 8.0, 2.4 Hz), 6.49 (d, 2H, $J$ = 8.4 Hz), 6.53 (s, 1H), 6.58 (d, 1H, $J$ = 2.4 Hz), 7.1-7.2 (m, 3H), 7.2-7.3 (m, 8H), 7.3-7.4 (m, 4H), 8.08 (d, 1H, $J$ = 2.0 Hz).

$^{13}$C NMR (100 MHz, DMSO-$d_6$): δ 1.0, 46.3, 70.9, 81.2, 83.8, 116.6, 117.5, 119.0, 120.1, 120.2, 125.3, 126.2, 127.0, 128.3, 129.4, 129.7, 130.8, 130.9, 131.1, 131.6, 132.2, 132.3, 138.1, 138.7, 143.8, 143.9, 144.0, 145.6, 145.7, 146.9, 148.1.

LRMS (ESI$^-$): $m/z$ = 755 ([M–H]$^+$)

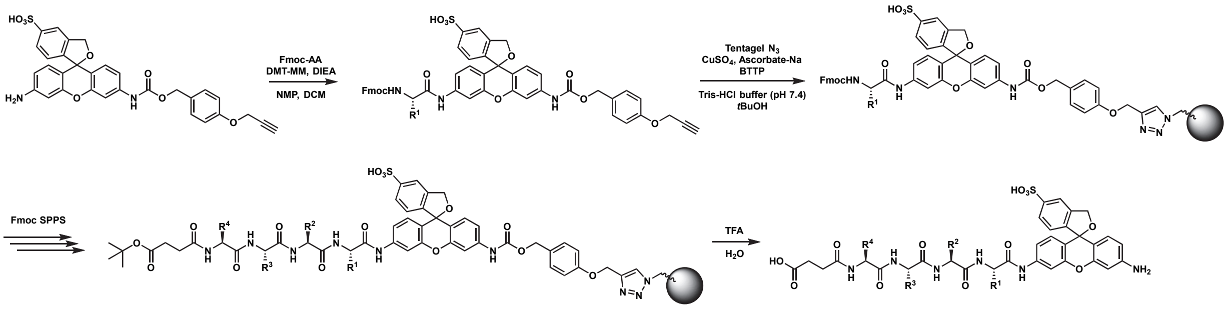


**Scheme S3.** Synthetic scheme of sHMRG-based probes

**Preparation of sHMRG-based probes**

All procedures were performed using an automated peptide synthesizer (Syro I, Biotage) equipped with a heating block.

(1) Amidation: sHMRG-Cpz (1 μmol) was mixed with Fmoc-protected amino acid building block (P1 amino acid, 10 μmol), DMT-MM (2.8 mg, 10 μmol) and DIEA (2.8 μL, 15 μmol) in NMP (10 μL) and DCM (500 μL), and the reaction was stirred at 65°C for 2 h.

(2) Capture and wash: Tentagel-azide beads (100 mg) were suspended in Tris-HCl buffer (300 mM, pH 7.4; 300 μL) in the reactor. Reaction mixture of (1) was diluted in $t$-BuOH (400 μL) and added into the reactor.

$CuSO_4$ (10 mM in $H_2O$, 100 µL), TBTA (30 mM in DMSO, 100 µL) and sodium ascorbate (30 mM in $H_2O$, 100 µL) were added to the reactor and stirred at 25°C for 2 h. The beads were washed six times with DMF.

(3) Peptide elongation: The peptide synthesis was performed by treating beads with conditions for (a) Fmoc deprotection and (b) amino acid coupling, successively until the full peptide sequence was prepared. (a) Piperidine (40% in DMF; 1200 µL) was added to beads and stirred at 25°C for 3 min. After removing the solvent, piperidine (40% in DMF; 600 µL) and DMF (600 µL) were added and stirred for 12 min. Beads were washed six times with DMF. (b) Fmoc-AA (0.4 M in DMF, 800 µL), HATU (0.4 M in DMF, 840 µL), DIEA (1.6 M in NMP, 400 µL) were added to beads and stirred at 30°C for 40 min. Beads were washed three times with DMF. Mono-*tert*-butyl succinate was used as a building block of Suc capping.

(4) Cleavage: 90% TFA *aq.* (200 µL) was added to the beads and stirred at 25°C for 15 min. The solution was collected, and the beads were washed three times with acetone (200 µL). The combined solution was diluted with $H_2O$ and freeze-dried. For the probes used for validation assay and probes with purity < 90% (500 nm), they were purified over prep. MPLC ($C_{18}$; $H_2O$-0.1% TFA/AcCN-0.1% TFA = 95/5 to 0/100 over 15 min). 10 mM DMSO stock solution was prepared by measuring the absorbance of compound in 0.1N HCl aq. and calculating the concentration based on ε of amide-protected HMRG to be 30,000[12].

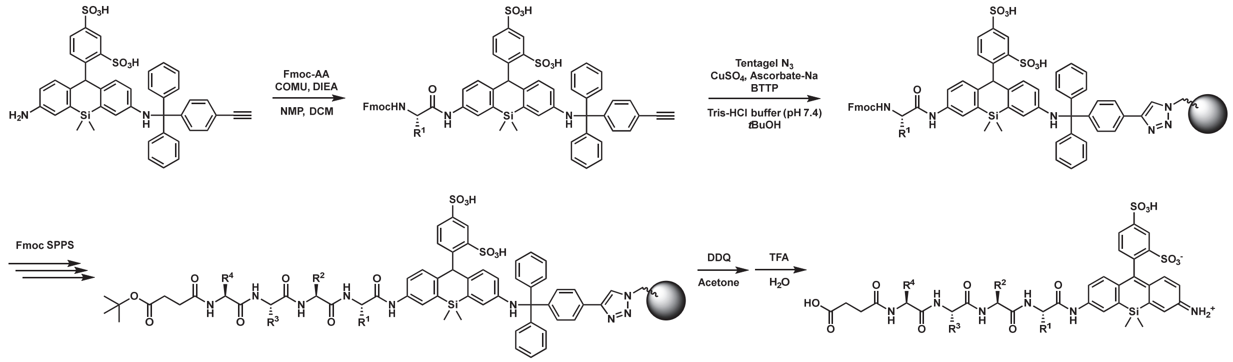


**Scheme S4.** Synthetic scheme of dsSiR-based probes

**Preparation of dsSiR-based probes**

All procedures were performed using an automated peptide synthesizer (Syro I, Biotage) equipped with a heating block.

(1) Amidation: leuco-dsSiR Ert (1 µmol) was mixed with Fmoc-protected amino acid building block (P1 amino acid, 10 µmol), COMU (4.3 mg, 10 µmol) and DIEA (5.6 µL, 30 µmol) in NMP (10 µL) and DCM (500 µL), and the reaction was stirred at 65°C for 2 h.

(2) Capture and wash: Tentagel-azide beads (100 mg) were suspended in Tris-HCl buffer (300 mM, pH 7.4; 300 µL) in the reactor. Reaction mixture of (1) was diluted in *t*-BuOH (400 µL) and added into the reactor. $CuSO_4$ (10 mM in $H_2O$, 100 µL), TBTA (30 mM in DMSO, 100 µL) and sodium ascorbate (30 mM in $H_2O$, 100 µL) were added to the reactor and stirred at 25°C for 2 h. The beads were washed six times with DMF.

(3) Peptide elongation: The peptide synthesis was performed by treating beads with conditions for (a) Fmoc deprotection and (b) amino acid coupling, successively until the full peptide sequence was prepared. (a) Piperidine (40% in DMF; 1200 μL) was added to beads and stirred at 25°C for 3 min. After removing the solvent, piperidine (40% in DMF; 600 μL) and DMF (600 μL) were added and stirred for 12 min. Beads were washed six times with DMF. (b) Fmoc-AA (0.4 M in DMF, 800 μL), HATU (0.4 M in DMF, 840 μL), DIEA (1.6 M in NMP, 400 μL) were added to beads and stirred at 30°C for 40 min. Beads were washed three times with DMF. Mono-*tert*-butyl succinate was used as a building block of Suc capping.

(4) Oxidation: 2,3-Dichloro-5,6-dicyano-1,4-benzoquinone (DDQ) (10 mg, 0.044 mmol) dissolved in acetone (1 mL) was added to the beads and stirred at 25°C for 3 h. Beads were washed six times with acetone.

(5) Cleavage: 90% TFA *aq.* (200 μL) was added to the beads and stirred at 25°C for 15 min. The solution was collected, and the beads were washed three times with acetone (200 μL). The combined solution was diluted with $H_2O$ and freeze-dried. For the probes used for validation assay and probes with purity < 90% (500 nm and 600 nm), they were purified over prep. MPLC ($C_{18}$; $H_2O$-0.1% TFA/AcCN-0.1% TFA = 95/5 to 0/100 over 15 min).

**Suc-AAVY-dsSiR**

LC Chromatogram was monitored at 500 nm ($H_2O$-0.1% TFA/AcCN-0.1% TFA = 95/5 to 0/100, 3.5 min).

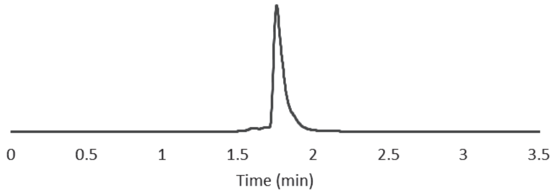


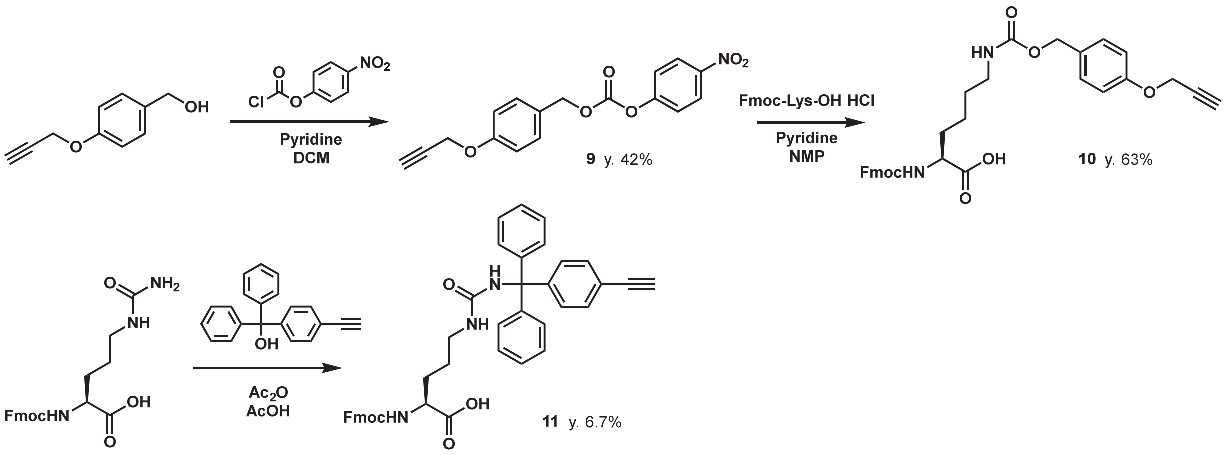


**Scheme S5.** Synthetic scheme of amino acid building blocks for SCCR.

**Preparation of Cpz-ONP (9)**

Pzl-OH (320 mg, 1.97 mmol) was dissolved in DCM (5 mL). 4-Nitrophenyl chloroformate (398 mg, 1.97 mmol) and pyridine (238 µL, 1.5 mmol) were added, and the reaction was stirred at 25°C for 3 h. The reaction was diluted with DCM, washed with 0.1N HCl *aq.* and brine, dried over $Na_2SO_4$, filtered, and evaporated. The product was purified over column chromatography (silica; Hexane/AcOEt = 100/0 to 0/100 over 15 min) to afford Cpz-ONP (270 mg, y. 42%) as white solid.

$^1$H-NMR (400 MHz, $CDCl_3$): δ 8.26 (d, 2H, *J* = 8.4 Hz), 7.42 (d, 2H, *J* = 8.4 Hz), 7.35 (d, 2H, *J* = 8.4 Hz), 7.00 (d, 2H, *J* = 8.4 Hz), 5.23 (s, 2H), 4.70 (s, 2H), 2.52 (s, 1H).

LRMS (ESI$^+$): *m/z* = 328 $(M+H)^+$

**Preparation of Fmoc-Lys(Cpz)-OH (10)**

Fmoc-Lys-OH HCl salt (80 mg, 0.20 mmol) was dissolved in *N*-methylpyrrolidone (NMP; 2 mL), and Cpz-ONP (13 mg, 0.04 mmol) and pyridine (48 µL, 0.59 mmol) were added, and the reaction was stirred at 25°C for 18 h. The reaction was diluted with DCM and subjected to silica gel column chromatography (silica; DCM/MeOH/AcOH = 99.9/0/0.1 to 89.9/10/0.1 over 15 min) to afford Fmoc-Lys(Cpz)-OH (28 mg, y. 63%) as white solid.

$^1$H-NMR (400 MHz, $CDCl_3$): δ 7.72 (d, 2H, *J* = 8.4 Hz), 7.56 (m, 2H), 7.35 (d, 2H, *J* = 8.4 Hz), 7.2-7.3 (m, 4H), 6.91 (m, 2H), 5.00 (s, 2H), 4.71 (s, 2H), 4.2-4.3 (m, 3H), 4.18 (t, 1H, *J* = 8.0 Hz), 3.2-3.3 (m, 2H), 2.48 (s, 1H), 1.2-1.8 (m, 6H).

LRMS (ESI$^+$): *m/z* = 557 $(M+H)^+$

**Preparation of Fmoc-Cit(Ert)-OH (11)**

Fmoc-Cit-OH (640 mg, 1.61 mmol) was dissolved in glacial acetic acid (1 mL). Ert-OH (307 mg, 1.08 mmol) and acetic anhydride (457 µL, 4.83 mmol) were added, and the reaction was stirred at 60°C for 30 min. Toluene was added to the reaction and the mixture was concentrated in vacuo. The remaining was diluted by AcCN and loaded directly into prep. MPLC for purification ($C_{18}$; $H_2O$-0.1% TFA/AcCN-0.1% TFA = 95/5 to 0/100 over 15 min). Freeze-drying of the fractions afforded Fmoc-Cit(Ert)-OH (48 mg, y. 6.7%) as white solid.

$^1$H-NMR (400 MHz, AcCN-$d_3$): δ 7.80 (d, 2H, *J* = 8.4 Hz), 7.63 (d, 2H, *J* = 8.4 Hz), 7.3-7.4 (m, 4H), 7.1-7.3 (m, 14H), 6.18 (s, 1H), 6.03 (2H, *J* = 8.0 Hz), 5.29 (br, 1H), 4.28 (d, 2H, *J* = 8.4 Hz), 4.21 (t, 1H, *J* = 8.4 Hz), 4.15 (m, 1H), 3.32 (s, 1H), 2.9-3.0 (m, 2H), 1.4-1.7 (m, 2H), 1.37 (m, 2H).

LRMS (ESI$^+$): *m/z* = 664 $(M+H)^+$

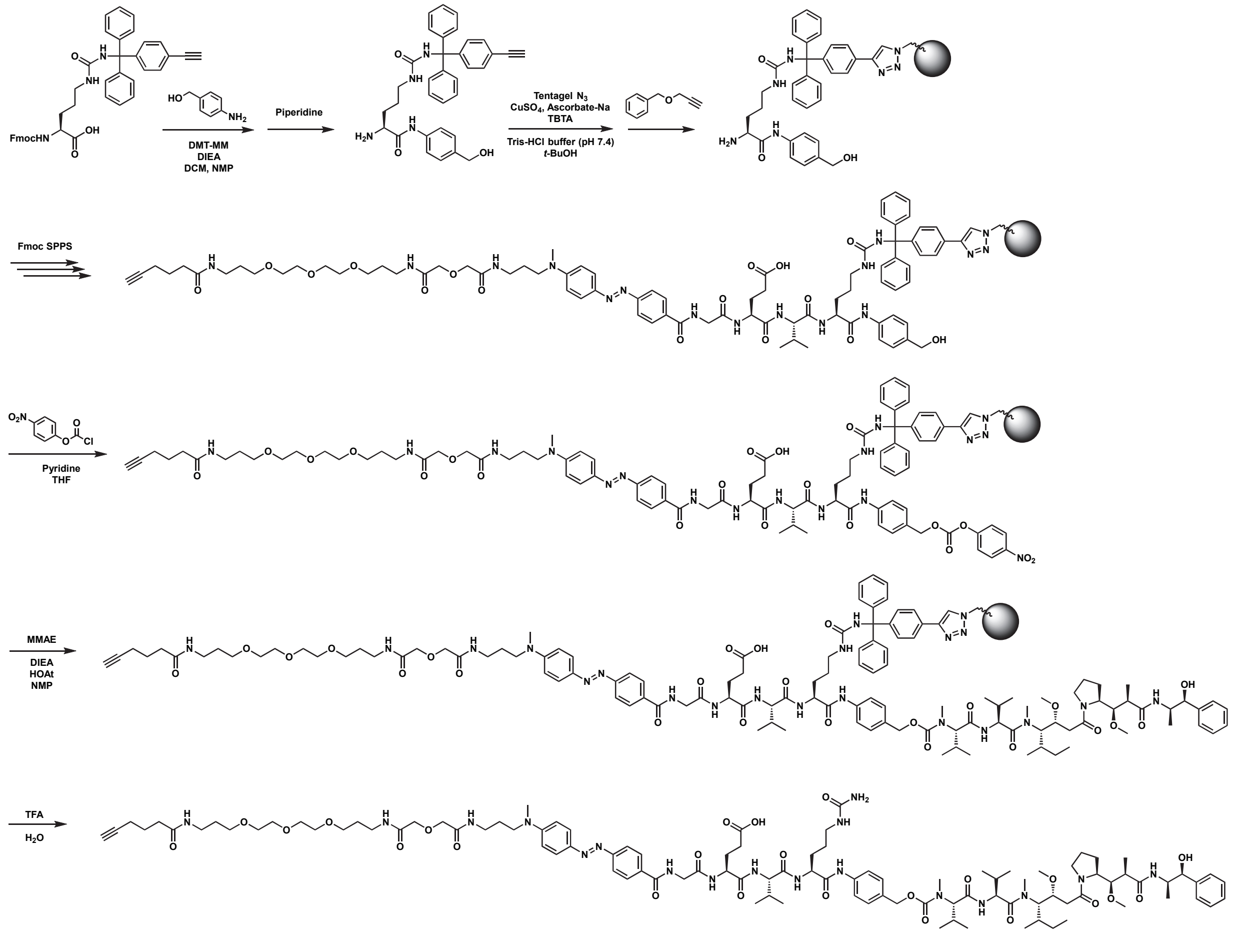

**Scheme S6.** Synthetic scheme of ADC linkers with SCCR.

### Preparation of ADC linkers

All procedures were performed using an automated peptide synthesizer (Syro I, Biotage) equipped with heating block kit.

(1) Amidation: Fmoc-Cit(Ert)-OH or Fmoc-Lys(Cpz)-OH (5 µmol) was mixed with *p*-aminobenzyl alcohol (6.5 mg, 50 µmol), DMT-MM (14.6 mg, 50 µmol) and DIEA (2.8 µL, 15 µmol) in NMP (10 µL) and DCM (500 µL), and the reaction was stirred at 65°C for 2 h.

(2) Capture and wash: Tentagel-azide beads (100 mg) were suspended in Tris-HCl buffer (300 mM, pH 7.4; 300 µL) in the reactor. Reaction mixture of (1) was diluted in *t*-BuOH (400 µL) and added into the reactor. $CuSO_4$ (10 mM in $H_2O$, 100 µL), TBTA (30 mM in DMSO, 100 µL) and sodium ascorbate (30 mM in $H_2O$, 100 µL) were added to the reactor and stirred at 25°C for 1 h. Benzyl propargyl ether (7.7 µL) in *t*-BuOH (100 µL) was added to the reactor and stirred at 25°C for 30 min. The beads were washed six times with DMF.

(3) Peptide elongation: The peptide synthesis was performed by treating beads with conditions for (a) Fmoc deprotection and (b) amino acid coupling, successively until the full peptide sequence was prepared. (a)

Piperidine (40% in DMF; 1200 μL) was added to beads and stirred at 25°C for 3 min. After removing the solvent, piperidine (40% in DMF; 600 μL) and DMF (600 μL) were added and stirred for 12 min. Beads were washed six times with DMF. (b) Fmoc-AA (0.4 M in DMF, 800 μL), DMT-MM (0.4 M in MeOH, 840 μL), DIEA (1.6 M in NMP, 400 μL) were added to beads and stirred at 30°C for 40 min. Beads were washed three times with DMF. Hexanoic acid was used as a building block of mAb modification site, Fmoc-NH-$PEG_2$-DGA-OH (Watanabe chemicals) was used as a building block of PEG linker, Fmoc-Apd-OH[13] was used as a building block of colorimetric reporter.

(4) 4-Nitrophenyl carbonate: The beads were washed six times with THF, and pyridine/THF (300 mM, 500 μL) was added. 4-Nitrophenyl chloroformate (10.6 mg, 53 μmol, 10×) was dissolved in THF (500 μL) and the solution was added to the beads. The reaction was stirred at 25°C for 1 h. The beads were washed six times with DMF.

(5) Attachment of payload: The beads were dissolved in DIEA (1.6 M in NMP, 50 μL), and MMAE or Sarcosine benzyl ester (7.5 μmol, 1.5×) in NMP (50 μL), HOAt (1.0 mg, 7.5 μmol) in NMP (50 μL) were added successively, and the reaction was stirred at 25°C for 12 h. The beads were washed six times with DMF and six times with THF. Use of higher amount of the payload for the increased concentration can increase the yield and purity of the product.

(6) Cleavage and purification: 90% TFA *aq.* (200 μL) was added to the beads and was stirred at 25°C for 5 min. The solution was collected, and beads were washed in three times with acetone (200 μL). The combined solution was diluted with $H_2O$ and was directly injected into prep. MPLC for purification ($C_{18}$; $H_2O$-0.1% TFA/AcCN-0.1% TFA = 95/5 to 0/100 over 15 min). Freeze-drying of the fractions afforded the desired product.

**Hex-PEG-Apd-GEVCit-PABC-MMAE (ADC #1).**

LC Chromatogram was monitored at 500 nm ($H_2O$-0.1% TFA/AcCN-0.1% TFA = 95/5 to 0/100, 3.5 min).

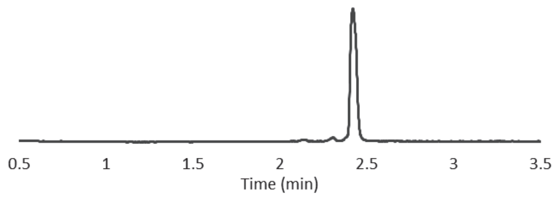


**Hex-PEG-Apd-GEVCit-PABC-Sar-OBzl (ADC #2)**

LC Chromatogram was monitored at 500 nm ($H_2O$-0.1% TFA/AcCN-0.1% TFA = 95/5 to 0/100, 3.5 min).

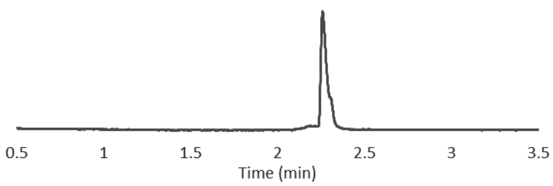

**Hex-Gly-Apd-EVCit-PABC-MMAE (ADC #3)**

LC Chromatogram was monitored at 500 nm ($H_2O$-0.1% TFA/AcCN-0.1% TFA = 95/5 to 0/100, 3.5 min).

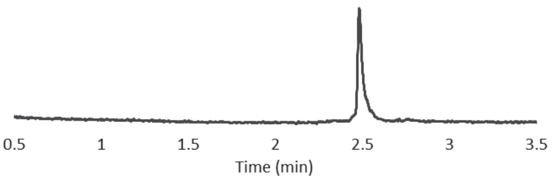


**Hex-PEG-Apd-AVCit-PABC-MMAE (ADC #4)**

LC Chromatogram was monitored at 500 nm ($H_2O$-0.1% TFA/AcCN-0.1% TFA = 95/5 to 0/100, 3.5 min).

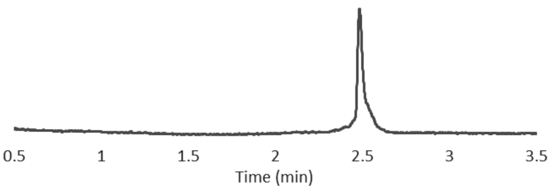


**Hex-PEG-Apd-GfFK-PABC-MMAE (ADC #5)**

LC Chromatogram was monitored at 500 nm ($H_2O$-0.1% TFA/AcCN-0.1% TFA = 95/5 to 0/100, 3.5 min).

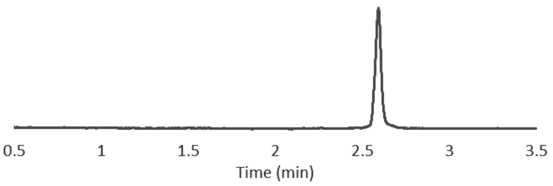


**Hex-PEG-Apd-GfFK-PABC-Sar-OBzl (ADC #6)**

LC Chromatogram was monitored at 500 nm ($H_2O$-0.1% TFA/AcCN-0.1% TFA = 95/5 to 0/100, 3.5 min).

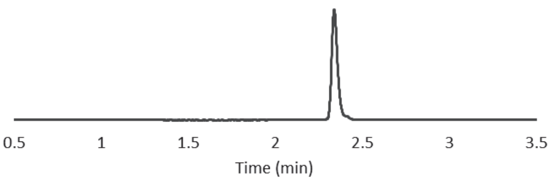

**Settings of automated peptide synthesizer and preparation of reagents**

**Reactions for preparation of sHMRG-based probes**

- Reaction
  - Reaction 2 h; 65°C; Vortex 1 min; Break 1 min
- Beads capture
  - Fill [a] 300 μL -> RV1
  - Fill [b] 200 μL -> RV2
  - Reaction 30 s; 25°C; Vortex 1 min; Break 1 min
  - Fill RV2 200 μL -> RV1
  - Fill [b] 200 μL -> RV2
  - Reaction 30 s; 25°C; Vortex 1 min; Break 1 min
  - Fill RV2 200 μL -> RV1
  - Fill [c] 100 μL -> RV1
  - Reaction 10 s; 25°C; Vortex 1 min; Break 1 min
  - Fill [d] 100 μL -> RV1
  - Reaction 10 s; 25°C; Vortex 1 min; Break 1 min
  - Fill [e] 100 μL -> RV1
  - Reaction 1 h; 25°C; Vortex 10 s; Break 1 min
  - Fill [f] 100 μL -> RV1
  - Reaction 30 min; 25°C; Vortex 10 s; Break 1 min
  - Empty 1 min
  - Wash 6 cycles
    - Fill [DMF] 1000 μL -> RV1
    - Reaction 30 s; 25°C; Vortex 10 s; Break 1 min
    - Empty 30 s
  - Fmoc deprotection
    - Fill [Piperidine/DMF] 1200 μL -> RV1
    - Reaction 3 min; 25°C; Vortex 10 s; Break 1 min
    - Empty 30 s
    - Fill [Piperidine/DMF] 600 μL -> RV1
    - Fill [DMF] 600 μL -> RV1
    - Reaction 12 min; 25°C; Vortex 10 s; Break 1min
    - Empty 10 s
- Amino acid elongation n cycles
  - Condensation
    - Fill [AA] 800 μL -> RV1
    - Fill [HATU/DMF] 840 μL -> RV
    - Fill [DIPEA/NMP] 400 μL -> RV
    - Reaction 40 min; 25°C; Vortex 15 s; Break 2 min
    - Empty 30 s
  - Fmoc deprotection
    - Fill [Piperidine/DMF] 1200 μL -> RV1
    - Reaction 3 min; 25°C; Vortex 10 s; Break 1 min
    - Empty 30 s
    - Fill [Piperidine/DMF] 600 μL -> RV1
    - Fill [DMF] 600 μL -> RV1
    - Reaction 12 min; 25°C; Vortex 10 s; Break 1 min
    - Empty 10 s

**Reagents for preparation of sHMRG-based probes**

| | | |
|---|---|---|
| RV1 | | |
| | Tentagel-azide | 100 mg |
| RV2 | | |
| | sHMRG-Cpz | 5 μmol |
| | Fmoc-AA | 10 μmol |
| | DMT-MM | 10 μmol |
| | DIEA | 2.8 μL |
| | NMP | 10 μL |
| | DCM | 500 μL |
| Amino acids | | |
| | [AA] | 0.4 M |
| System liquids | | |
| | [Piperidine/DMF] | 40% (v/v) |
| | [DMT-MM/MeOH] | 0.4 M |
| | [DIPEA/NMP] | 1.6 M |
| Reagents | | |
| [a] | Tris-HCl buffer (pH 7.4) | 300 mM |
| [b] | *t*-BuOH | |
| [c] | $CuSO_4/H_2O$ | 10 mM |
| [d] | TBTA/DMSO | 30 mM |
| [e] | Sodium ascorbate/$H_2O$ | 30 mM |

**Reactions for preparation of ADC linkers**

Reaction

Reaction 2 h; 65°C; Vortex 1 min; Break 1 min

Beads capture

Fill [a] 300 μL -> RV1

Fill [b] 200 μL -> RV2

Reaction 30 s; 25°C; Vortex 1 min; Break 1 min

Fill RV2 200 μL -> RV1

Fill [b] 200 μL -> RV2

Reaction 30 s; 25°C; Vortex 1 min; Break 1 min

Fill RV2 200 μL -> RV1

Fill [c] 100 μL -> RV1

Reaction 10 s; 25°C; Vortex 1 min; Break 1 min

Fill [d] 100 μL -> RV1

Reaction 10 s; 25°C; Vortex 1 min; Break 1 min

Fill [e] 100 μL -> RV1

Reaction 1 h; 25°C; Vortex 10 s; Break 1 min

Fill [f] 100 μL -> RV1

Reaction 30 min; 25°C; Vortex 10 s; Break 1 min

Empty 1 min

Wash 6 cycles

Fill [DMF] 1000 μL -> RV1

Reaction 30 s; 25°C; Vortex 10 s; Break 1 min

Empty 30 s

Fmoc deprotection

Fill [Piperidine/DMF] 1200 μL -> RV1

Reaction 3 min; 25°C; Vortex 10 s; Break 1 min

Empty 30 s

Fill [Piperidine/DMF] 600 μL -> RV1

Fill [DMF] 600 μL -> RV1

Reaction 12 min; 25°C; Vortex 10 s; Break 1min

Empty 10 s

Amino acid elongation n cycles

Condensation

Fill [AA] 800 μL -> RV1

Fill [HATU/DMF] 840 μL -> RV

Fill [DIPEA/NMP] 400 μL -> RV

Reaction 40 min; 25°C; Vortex 15 s; Break 2 min

Empty 30 s

Fmoc deprotection

Fill [Piperidine/DMF] 1200 μL -> RV1

Reaction 3 min; 25°C; Vortex 10 s; Break 1 min

Empty 30 s

Fill [Piperidine/DMF] 600 μL -> RV1

Fill [DMF] 600 μL -> RV1

Reaction 12 min; 25°C; Vortex 10 s; Break 1 min

Empty 10 s

Capping

Fill [DMF] 200 μL -> RV1

Fill [DIPEA/NMP] 200 μL -> RV1
Fill [g] 100 μL -> RV1
Reaction 1 h; 25°C; Vortex 10 s; Break 1 min
Empty 1 min
Wash 3 cycles
Fill [DMF] 2000 μL -> RV1
Reaction 30 s; 25°C; Vortex 10 s; Break 1 min
Empty 30 s

*p*-Nitrophenyl carbonate formation
Wash 6 cycles
Fill [THF] 2000 μL -> RV1
Reaction 30 s; 25°C; Vortex 10 s; Break 1 min
Empty 30 s
Fill [Pyridine/THF] 500 μL -> RV1
Fill [THF] 1000 μL -> [h]
Fill [h] 500 μL -> RV1
Reaction 30 min; 25°C; Vortex 10 s; Break 1 min
Empty 1 min
Wash 6 cycles
Fill [DMF] 2000 μL -> RV1
Reaction 30 s; 25°C; Vortex 10 s; Break 1 min
Empty 30 s

Payload loading
Fill [DIPEA/NMP] 50 μL -> RV1
Fill [i] 50 μL -> RV1
Fill [j] 50 μL -> RV1
Reaction 12 h; 25°C; Vortex 10 s; Break 1 min
Empty 1 min
Wash 3 cycles
Fill [DMF] 2000 μL -> RV1
Reaction 30 s; 25°C; Vortex 10 s; Break 1 min
Empty 30 s
Wash 6 cycles
Fill [THF] 2000 μL -> RV1
Reaction 30 s; 25°C; Vortex 10 s; Break 1 min
Empty 30 s

**Reagents for preparation of ADC linkers**

| | | |
|---|---|---|
| RV1 | | |
| | Tentagel-azide | 100 mg |
| RV2 | | |
| | SCCR-compatible amino acid building blocks | 5μmol |
| | *p*-Aminobenzyl alcohol | 50 μmol |
| | DMT-MM | 50 μmol |
| | DIEA | 2.8 μL |
| | NMP | 10 μL |
| | DCM | 500 μL |
| Amino acids | | |
| | [AA] | 0.4 M |
| System liquids | | |
| | [Piperidine/DMF] | 40% (v/v) |
| | [DMT-MM/MeOH] | 0.4 M |
| | [DIPEA/NMP] | 1.6 M |
| | [Pyridine/THF] | 300 mM |
| Reagents | | |
| [a] | Tris-HCl buffer (pH 7.4) | 300 mM |
| [b] | *t*-BuOH | |
| [c] | $CuSO_4/H_2O$ | 10 mM |
| [d] | TBTA/DMSO | 30 mM |
| [e] | Sodium ascorbate/$H_2O$ | 30 mM |
| [f] | Benzyl propargyl ether/*t*-BuOH | 500 mM |
| [g] | Capping reagents/NMP | 100 mM |
| [h] | 4-Nitrophenyl chloroformate | 20 mg |
| [i] | Payload/NMP | 150 mM |
| [j] | HOAt/NMP | 150 mM |
